# The antifungal capacity of a 681-membered collection of environmental yeast isolates

**DOI:** 10.1101/2024.07.29.605670

**Authors:** Alicia Maciá Valero, Fatemehalsadat Tabatabaeifar, Sonja Billerbeck

**Affiliations:** Department for Molecular Microbiology, University of Groningen, Nijenborgh 7, 9747 AG Groningen, the Netherlands

**Keywords:** bioprospecting, novel antifungals, killer yeast, screening, iron siderophores

## Abstract

Fungal pathogens threaten human health and food security, with resistance reported across limited antifungal classes. Novel strategies to control these pathogens and food spoilers are urgently needed.

Environmental yeasts provide a functionally diverse, yet underexploited potential for fungal control based on their natural competition via the secretion of iron siderophores, killer toxins (proteins) or other small molecules like volatile organic compounds or biosurfactants. However, there is a lack of standardized workflows to systematically access application-relevant yeast-based compounds and understand their molecular functioning.

Towards this goal, we developed a workflow to identify and characterize yeast isolates that are active against relevant human and plant pathogens and spoilage yeasts, herein focusing on discovering yeasts that produce potential killer toxins. The workflow includes the classification of the secreted molecules and cross-comparison of their antifungal capacity using an independent calibrant.

Our workflow delivered a collection of 681 yeasts of which 212 isolates (31%) displayed antagonism against at least one of our target strains. While 50% of the active yeasts showed iron-depended antagonism, likely due to siderophore production, more than 25% are potentially secreting a toxic protein. Those killer yeast candidates clustered within ten species, showed target profiles from narrow- to broad spectrum, and several showed a broad pH and temperature activity profile.

Given the tools for yeast biotechnology and protein engineering available, our collection offers a foundation for genetic and molecular characterization of antifungal phenotypes, with potential for future exploitation. The scalable workflow can screen other yeast collections or adjust for different antifungal compounds.

## Introduction

Fungal pathogens infect plants and humans and thus pose an urgent threat to society. A third of total crops produced worldwide is lost due to different species of fungi .^1,2^ Moreover, 25% of the spoilage in post-harvest fruits, dairy products or other processed foods is caused by microbial contamination (up to 30% in North America).^3^ In addition, fungi are responsible for the spoilage of fermented beverages, leading to significant economic losses every year.^4^ Some examples of fungal pathogens are *Botrytis cinerea, Penicillium digitatum* and *Geotrichum candidum* in crops,^2,5^ *Zygosaccharomyces* spp and *Penicillium* spp in food^5–7^ or *Brettanomyces* in the fermented drinks industry.^8^

Further an estimated 3.8 million people pass annually from fungal infections^9^. Fungal diseases constitute a specifically high burden to individuals with a compromised immune system, such as HIV-positives, chemotherapy patients or individual with a COVID-19 infection^10–12^. The clinically most relevant species are *Candida albicans, Nakasomyces glabratus* (previously known as *Candida glabrata*)*, Candida tropicalis, Candida parapsilosis* and *Candida krusei*^13,14^, causing more than 135 million cases of candidiasis^15^ and more than 1.4 million cases of candidemia^9^ around the world. Recently, *Candida auris* has gained public attention due to the emergence of multi-resistant isolates in several countries around the globe, including the Netherlands^16–20^, and it was classified in 2019 as urgent antibiotic resistant threat by the Center for Disease Control and Prevention in the United States (CDC)^21^. Subsequently, the World Health Organization published their first list of health-threatening fungi in 2022, where *C. auris* and *C. albicans* are regarded as critical priority pathogens and *N. glabratus, C. parapsilosis*and *C. tropicalis* are listed in the high priority group^22^.

There are mainly four classes of antimicrobials to treat fungal infections in humans: the azoles, the echinocandins, the polyenes and the pyrimidine analogues. However, resistance to all classes has been reported in different fungal pathogens, including the most prevalent *Candida* species.^23–30^

The azoles are not only used in the clinic but are also the preferred antifungals in agriculture ^31^. While the widespread application of azole-based pesticides has historically proven highly effective in controlling fungal pest, their excessive usage, coupled with crop monoculturing, temperatures rising and intensified global trade, has facilitated the worldwide dissemination of fungal diseases and led to a sharp emergence in resistance^32^. In addition, the European Commission has enforced halving the use of chemical pesticides in agriculture by 2030.^33^

For the treatment of human pathogens, amphotericin B (polyene) is the gold standard in clinical use. However, its activity is often inefficient, requiring a combination of different antifungal classes,^31^ prolonged exposure or higher doses leading to toxicity.^34^

Thus, there is an urgent need for new antifungals, both for agriculture and human health.^35^ Ideally, these alternatives should possess diverse modes of action and target ranges allowing for a decrease in the simultaneous utilization of identical molecule classes across agriculture, food processing and medical settings and therefore decreasing the possibility of resistance emerging.^36^ However, the development of novel antifungals is challenging, due to the limited cellular targets that distinguish the eukaryotic host and the pathogen.^37^

Yeasts are known to synthesize different compounds that antagonize fungi. Some yeast have already been explored as products for biocontrol.^38–40^ Antifungal compounds secreted by yeast include siderophores that capture environmental iron (and therefore deplete antagonists from essential nutrients) and other small molecules with unstudied modes of action such as biosurfactants and volatile organic compounds. Most prominently, yeast are known for secreting antifungal proteins which can either be enzymes or toxins that kill other fungi, the latter known as yeast killer toxins.

Yeast killer toxins have a unique potential to serve as templates for antifungal development as they represent a potentially vast but largely unexplored pool of functional diversity. From the few toxins that have been genetically identified and characterized, it is known that they present diverse unrelated sequences, cellular targets and toxicity mechanisms, ranging from pore formation, inhibition of cell-wall biosynthesis, cell-cycle arrest or induction of apoptosis to tRNAse activity^41–44^. While yeast killer toxins point us to new, evolutionary and optimized ways to target fungi, most known toxins have inherent limitations such as a narrow target range and a narrow activity profile restricting them to be only functional at low pH and low temperature.^36^ Nevertheless, some yeast killer toxins act in a broad spectrum^45,46^ and across a broad pH and temperature range.^47,48^ It is theorized that the majority of killer yeast diversity remains undiscovered.^36^ This diversity might include more toxins with broader application- friendly characteristics.

While protein engineering could be applied to improve the pH range and thermostability of yeast killer toxins, ideally such efforts are templated by a toxin that already shows a target range and physico-chemical properties that match as closely as possible an envisioned application. As such, there is a need for systematically mapping the yeast toxicome and its performance spectrum in terms of target range and physico-chemical requirements.

Bioprospecting for killer yeast has been performed in numerous occasions by isolating yeasts and screening them for activity against different yeast and filamentous fungi.^49–55^ However, the frequency of a true killer phenotype – based on protein secretion – is difficult to estimate from these approaches, as they often exclusively rely on simple halo assay readouts and do not discriminate killer strains that secrete a protein toxin from isolates that secrete other antifungal compounds.^51–57^ Further, systematic mapping of the target range against application-relevant yeasts and filamentous fungi as well as mapping of physico-chemical properties required for activity of the antifungal phenotype is often absent.

In this work, we developed and applied a systematic workflow that can access yeasts from the environment and provides first insights into the molecule classes causing this antimicrobial behaviour. The workflow provides an estimate on the prevalence of true killer yeasts and shows that killer yeasts with diverse target ranges from narrow to broad and different pH and temperature are virtually found in every niche. Future efforts on the genetic and molecular characterization of these yeast isolates and their toxins will open their exploitation on diverse applications, *e. g.*, pre- and post-harvest biocontrol agents, fermentation starters and biopharmaceutical producing platforms.

## Materials and Methods

### Materials, equipment and services

Isopropanol was obtained from Biosolve BV (Valkenswaard, The Netherlands). All remaining media and buffer components were obtained from BD Bioscience (Franklin Lakes, NJ, USA), Merck KGaA (Darmstadt, Germany) and Thermo Fisher Scientific Inc (Waltham, MA, USA).

Phire Hot Start II DNA polymerase and the Pico Green Kit were obtained from Thermo Fisher Scientific Inc (Waltham, MA, USA). Primers were obtained from Biolegio (Nijmegen, the Netherlands). Sterile tubes and round Petri dishes were obtained from Sarstedt AG & Co (Nümbrecht, Germany). Sterile one-well plates and round-bottom 96-well microtiter plates were obtained from Greiner Bio-One BV (Alphen aan den Rijn, the Netherlands). HTS Transwell®-96 permeable sterile membranes for 96-well plates were obtained from Corning (Corning, NY, USA). Centrifugal units Amicon® Ultra-4 3KDa molecular weight cut-off (MWCO) were obtained from Merck KGaA (Darmstadt, Germany) and 0.2 μm pore size Whatman™ filters, from Cytiva (Marlborough, MA, USA).

The manual pinner was a VP 409 MULTI-BLOT Replicator® obtained from V&P Scientific Inc (San Diego, CA, USA). The microscope used in this study was a Nikon Ti-E microscope from Nikon Instruments (Amstelveen, the Netherlands) equipped with a Hamamatsu Orca Flash4.0 camera. Images of screening plates for halo size measurement were obtained with a Fujifilm LAS4000 imager from GE Healthcare (Chicago, IL, USA).

Sanger sequencing services were provided by Macrogen Europe (Amsterdam, the Netherlands) and PacBio Next Generation Sequencing was performed by Biomarker Technologies (BMK) GmbH (Münster, Germany).

### Environmental sampling and yeast isolation

Environmental samples were collected from natural sources at different locations across the Netherlands (see **Supplementary Table 1)**. Samples were collected in sterile plastic tubes and either processed immediately or stored at 4 °C for up to one week. According to the substrate, they were classified as *water* (including sea, lakes, rivers, puddles and canals), *soil* (soil and sand), *plant material* (including flowers, trunk trees, leaves and fruits) and *other* (animal faeces and mushrooms).

Sample material was inoculated in shake flasks containing 20 mL of Yeast extract Peptone Dextrose medium (YPD: 2.5 g/L yeast extract, 5 g/L peptone, 20 g/L glucose) buffered to pH 4.0±0.1 (following the 0.2x scheme of the citrate-phosphate buffer system^58^) and containing ampicillin (100 μg/mL). YPD broth used for enrichments after October 2021 also contained 20 µg/mL FeCl_3._ Cultures were incubated for 48-72 hours. Incubation was carried out at 25 °C for samples collected in autumn/winter and 30 °C for samples acquired during spring/summer (see **Supplementary Table 1**). Following enrichment, three rounds of single colony isolation were performed by streaking on YPD agar plates containing ampicillin (100 μg/mL), spectinomycin (50 μg/mL) or kanamycin (50 μg/mL). Plates were incubated at 30 °C for 24 – 48 h. Single colonies of morphologically distinct yeasts were picked and analysed under the microscope at 100x magnification using 1% (w/v) agarose in phosphate- buffered saline pads to verify the purity of those yeast-like cultures. In order to array the entire collection into a 96-well format, each yeast isolate was inoculated into one well of a total of eight 96-well plates containing YPD broth with 10 % (v/v) glycerol. Yeasts were allowed to grow at 30 °C for 24 hours shaking. These plates were replicated on YPD agar one-well plates with a manual pinner and incubated at 30 °C. For further screening, the one-well plates were stored at 4 °C (source plates); for long-term storage, the 96-well plates were sealed with permeable sterile membranes and stored at -80 °C.

### Target strains and growth conditions

All target and control strains used in this study are listed in **Supplementary Table 2**. All microorganisms were grown on YPD agar medium at 30 °C for storage and in our in-house optimized synthetic complete (SC/Urea) medium^58^ at 25 °C when tested for antifungal activity, unless stated differently. Every media used in this study was buffered to pH 4.0±0.1 with 40 mM Na_2_HPO_4_ and 30 mM citric acid following the 0.2x scheme of the citrate-phosphate buffer system^58^ unless otherwise specified.

### Pooled species identification of the yeast collection via PacBio sequencing

The collection of yeast isolates previously arrayed in 96-well plates was replicated into fresh 96-well plates with 200 µL YPD broth and incubated at 30 °C shaking for 48 hours. Yeast cultures were pooled per plate by pipetting the volume of each well into a cell culture reservoir with a multi-channel pipette and transferring the combined cultures into a Falcon tube for further processing; this resulted in eight samples. Pooling per plate rather than pooling the entire collection was performed to reduce the amount of distinct yeast isolates present in each PacBio sequencing sample and hence increase the resolution and sensitivity of the sequencing results.

DNA isolation was performed based on a protocol adapted from de Kloet and colleagues^59^: the eight yeast samples were pelleted by centrifugation. The resulting pellets were re- suspended and incubated with 200U lyticase in 5 mL 50 mM EDTA-Na pH 8.0±0.1 buffer on a rolling platform at room temperature overnight. Subsequently, cells were pelleted, re- suspended in 4 mL of 2% (w/v) SDS and 50 mM EDTA-Na pH 8.0 buffer and incubated at 65 °C for 1 hour. Then, 1 mL 3 M potassium acetate was added and samples were mixed by vortexing. Cell debris were centrifuged for 10 minutes and the supernatants were transferred to new microcentrifuge tubes, where equal volumes of isopropanol were added. After mixing properly, samples were centrifuged to precipitate the DNA. The resulting pellet was washed with 70% (v/v) ethanol, dried and re-suspended in warm milliQ water (60 – 65 °C). DNA concentration was measured using a Quant-iT PicoGreen dsDNA assay kit and the eight different samples were outsourced for PacBio Next Generation Sequencing.

The company amplified the internal transcribed spacers (ITS) of the eight samples using primers ITS1 and ITS4 (**Supplementary Table 3**). The amplicons obtained were sequenced by the company and reads with high quality were subjected to computational length filtration, chimera removal and BLAST using the UNITE database. Eventually our lab was provided with a list of Amplicon Sequence Variants (ASVs). Those were blasted using standard databases and ascribed to the most closely related species based on sequence similarity and query coverage (see **Supplementary Table 4**). Afterwards, ASVs linked to the same species were clustered into Operational Taxonomic Units (OTUs).

### Species identification of individual isolates by Sanger sequencing

Yeasts of interest were species identified amplifying the conserved regions ITS1 region or D1/D2 domain using primer pairs P-ITS1 and ITS2, and NL1 and NL4, respectively (**Supplementary Table 3)**. The amplicons were cleaned enzymatically using ExoSAP-IT and Sanger sequenced. The sequences were blasted and the hits with highest percentage of sequence similarity and query coverage were the species assigned to each yeast sample.

### Antifungal activity screening assay

The eight source plates containing the wild yeast collection were pinned into 96-well plates pre-filled with 80 µL sterile milliQ water in order to create a cell suspension. From there, cells were replicated onto SC medium agar screening plates. The screening plates contained 0.003% (w/v) methylene blue^60^ and a target yeast strain seeded at 0.01 final OD_600_ units. Screening plates were incubated at 25 °C for 48 hours (72 hours for *B. bruxellensis*). Every isolate was tested against the eight yeast target strains shown in **Supplementary Table 2** in triplicates.

### Iron-depletion assay

All yeast isolates that had shown a growth inhibition halo against one or more target strains were arrayed in three new 96-well plates and screened following the method described above using plates supplemented with and without 20 µg/mL FeCl_3_. Every plate was screened three times.

### Assay for protein content in the spent culture media

All yeast isolates displaying iron-independent antifungal activity were grown individually in 5 mL SC medium glass culture tubes at 25 °C shaking for 24 hours. Cells were pelleted and stored at 4 °C. The spent culture media of each sample was filtered using a 0.2 μm pore size filter membrane. The obtained cell-free spent culture was concentrated 10 – 40 – fold using centrifugal units with a 3 KDa MWCO. For each yeast, 5 µL of the pelleted cells and 10 µL of concentrated spent culture media were assayed for antifungal activity against *C. tropicalis, S. cerevisiae* and *B. bruxellensis*. The screening plates were incubated at 25 °C for 24 hours.

### Measurement of antifungal halo sizes and calibration using a chemical standard

The killer yeast candidates producing antifungal proteins were re-arrayed in a new 96-well plate and screened as previously described against the eight target strains listed in **Supplementary Table 2.** Every plate included 5 µL spots of 8, 4, 2, 1 and 0.5 µg/mL micafungin sodium dissolved in milliQ water. Plates were incubated at 25 °C for 48 hours (72 hours for *B. bruxellensis*) and the halos formed by both killer yeast candidates and micafungin were measured in arbitrary units using ImageJ software. This experiment was performed in triplicates. Measurements of the halo areas formed at different micafungin concentrations were plotted over the concentration and fitted to a logarithmic function. We determined the equation of this function and used it to convert the halo area measurements of killer yeasts candidates into micafungin equivalents.

### Antifungal activity under different pH values and temperatures

The spent culture media of isolates yAMV76, yAMV92, yAMV116, yAMV254, yAMV368, yAMV382, yAMV384, yAMV499, yAMV598 and yAMV650 was 100 – fold concentrated as previously described to test the activity under different pH and temperature values. In short, these yeasts were grown in 100 mL SC medium at 25 °C for 24 hours. The cells were pelleted and their spent culture media was filtered. The resulting cell-free samples were concentrated 100 – fold using 3 KDa MWCO centrifugal units. The obtained concentrate was tested for antifungal activity by spotting 10 µL on SC medium agar plates seeded at 0.01 OD OD_600_ final units of *C. tropicalis.* Six different conditions were tested: SC media buffered to pH 4.0±0.1 and incubated at 25 °C, 30 °C and 37 °C; and SC media buffered to pH 7.0±0.1 (buffered with 80 mM Na_2_HPO_4_ and 8 mM citric acid) and incubated at 25 °C, 30 °C and 37 °C. All plates were incubated for 48 hours. This experiment was performed three times. Results were imaged and halos were measured in arbitrary units using ImageJ software.

### Effect on mycelial growth and conidiation in B. cinerea

The effect on mycelial growth and conidia formation in *B. cinerea* was determined as follows: First, a suspension of *B. cinerea* spores was inoculated and grown on Yeast Mannitor Agar plates (YMA; 1 g/L yeast extract, 10 g/L mannitol, 20 g/L agar, 0.2 g/L K_2_HPO_4_, 0.2 g/L MgSO_4_, 0.1g/L NaCl) buffered to pH 4.0±0.1 at 20 °C for 7 days. Then, an 8.5 mm mycelial agar plug was cut and placed in the centre of fresh YMA plates and incubated at 20 °C. The following day, every killer yeast candidate was individually inoculated into SC/urea media buffered to pH 4.0±0.1 and grown at 25 °C shaking for 24 hours. Afterwards, 1 mL of each culture was pelleted, and re-suspended in 50 µL, from which 10 µL of that cell suspension were spotted on three positions surrounding the mycelial plug. Plates were incubated at 20 °C and photos were taken after seven days.

For yAMV116, yAMV368 and yAMV598, the experiment was also conducted using 5 µL 200 – fold spent culture media, prepared as previously described.

### Software

All halo area measurements were performed using ImageJ 1.53u (rsb.info.nih.gov/ij/)^61^.

Every heatmap present in this manuscript was generated using GraphPad Prism® v9.1 software^62^ (GraphPad Software, Inc., 2013).

Multiple sequence alignments were performed using Clustal Omega Tool of the EMBL Job Dispatcher^63^ and visualised with Jalview 10.0.5^64^.

The phylogenetic tree was generated using the Maximum Likelihood method of W-IQ-TREE Web server^65^ (http://iqtree.cibiv.univie.ac.at/) after 500 iterations. The robustness of the branches was assessed with 1000 bootstrap replications.

## Results

### A systematic workflow for bioprospecting yeasts with antifungal activity

Our workflow starts with isolating yeasts from the environment (**Figure 1A**), screening those against a panel of eight relevant target fungi, followed by a screen yielding insights into the molecule classes they produce (**Figure 1B**). Our assays can discriminate between iron siderophores, proteins or neither. The workflow includes the pooled species identification of yeast isolates present in the collection and individually of those of interest (**Figure 1C**). Focussing herein on killer yeast candidates – meaning those yeast potentially secreting an active protein – we compared the activity range of isolates from the same species and identified profiles of potentially different proteins (**Figure 1D**). Further tested their pH and temperature activity ranges (**Figures 1E**).

**Figure 1.**
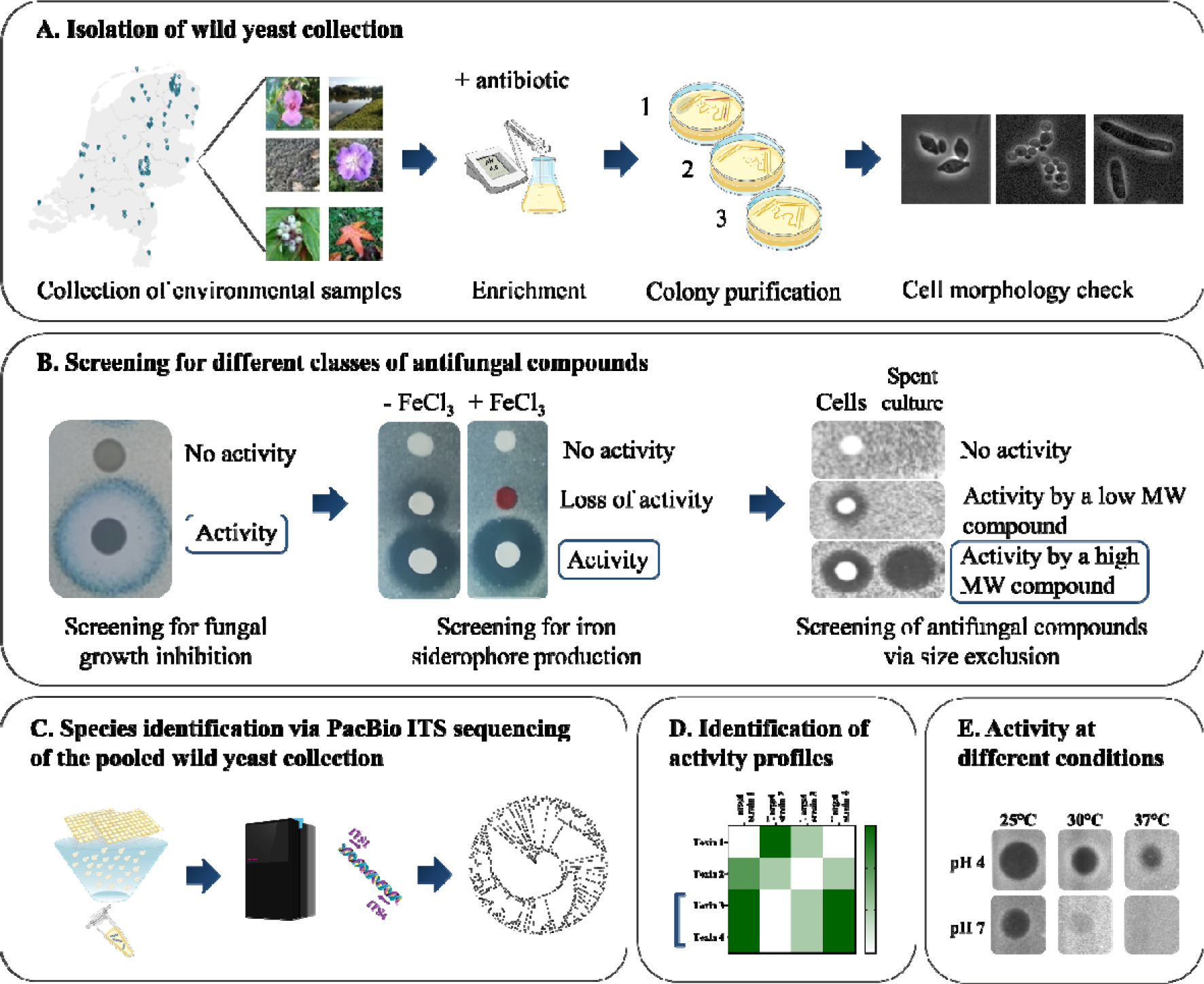
Workflow for isolating yeast-based antifungals from environmental samples. **A.** Isolation of wild yeasts. **B.** Screening for high molecular weight compounds with antifungal activity. **C.** Species identification of the yeast collection via pooling, DNA extraction and PacBio sequencing of the PCR-amplified ITS regions. **D**. Identification of activity profiles within the isolates. **E**. Test for activity under different temperature and pH conditions of spent media.

Our workflow does not yet include the genetic and molecular identification of the secreted molecules. Our panel of target fungi is composed of eight yeast species and one filamentous fungus representing some of the most notorious human pathogens, food spoilers and plant pests **(Supplementary Table 2).**

We considered design principles for scalability (*e.g.* for automation in bio-foundries) and cross-comparability such that more candidate yeasts can be screened in the future (*e.g.* those already available in strain collections) and such that results can be compared across studies and laboratories. First, we used single-celled target fungi (yeast) as the starting set for screening, as those grow in suspension and allow for high-throughput compatible antifungal assays. Those yeasts that presented an antagonistic phenotype, can then be subjected to lower throughput screens against filamentous fungi of interest.

We suggest a set of target fungi that are available in public strain collections and we developed a protocol to calibrate the antifungal capacity of the yeast to a commercially available antifungal (micafungin) and provided the activity in calibrated units.

Several technical considerations, such as false negatives due to specific screening conditions and the detection limit of the screen, are discussed within the conclusion.

### Biodiversity of the 681-membered wild yeast collection from the Netherlands

1,314 environmental samples were collected and classified depending on their substrate as *water* (from sea, lakes, rivers, puddles and canals), *soil* (soil and sand), *plant material* (including flowers, trunk trees, leaves and fruits) and *other* (animal faeces and mushrooms). From those samples, 681 different microorganisms were isolated and defined as yeasts (**Supplementary Table 1**), based on their ability to grow at low pH (4.0 – 4.5) and in the presence of any of the three antibiotics tested (spectinomycin, ampicillin or kanamycin), their unicellular growth on agar plate and their cellular morphology under the microscope. With the objective of maximizing species diversity and antifungal capacity, we incorporated yeasts that showed reproduction via budding and fission (**Figure 1A**), as well as yeasts presenting dimorphic growth^66^ (*i.e.,* able to grow as yeast and mycelium depending on the growth conditions), given that all of those had been reported to produce antifungal compounds before.^51,52,57,67,68^

From these 681 isolates, 533 (78%) were isolated from plant material, 79 (12%) from soil, 64 (9%) from water samples and five (1%) from other substrates (**Figure 2A**). The 681 yeast isolates were arrayed in a 96-well format (8 plates in total) for further handling and storage.

**Figure 2.**
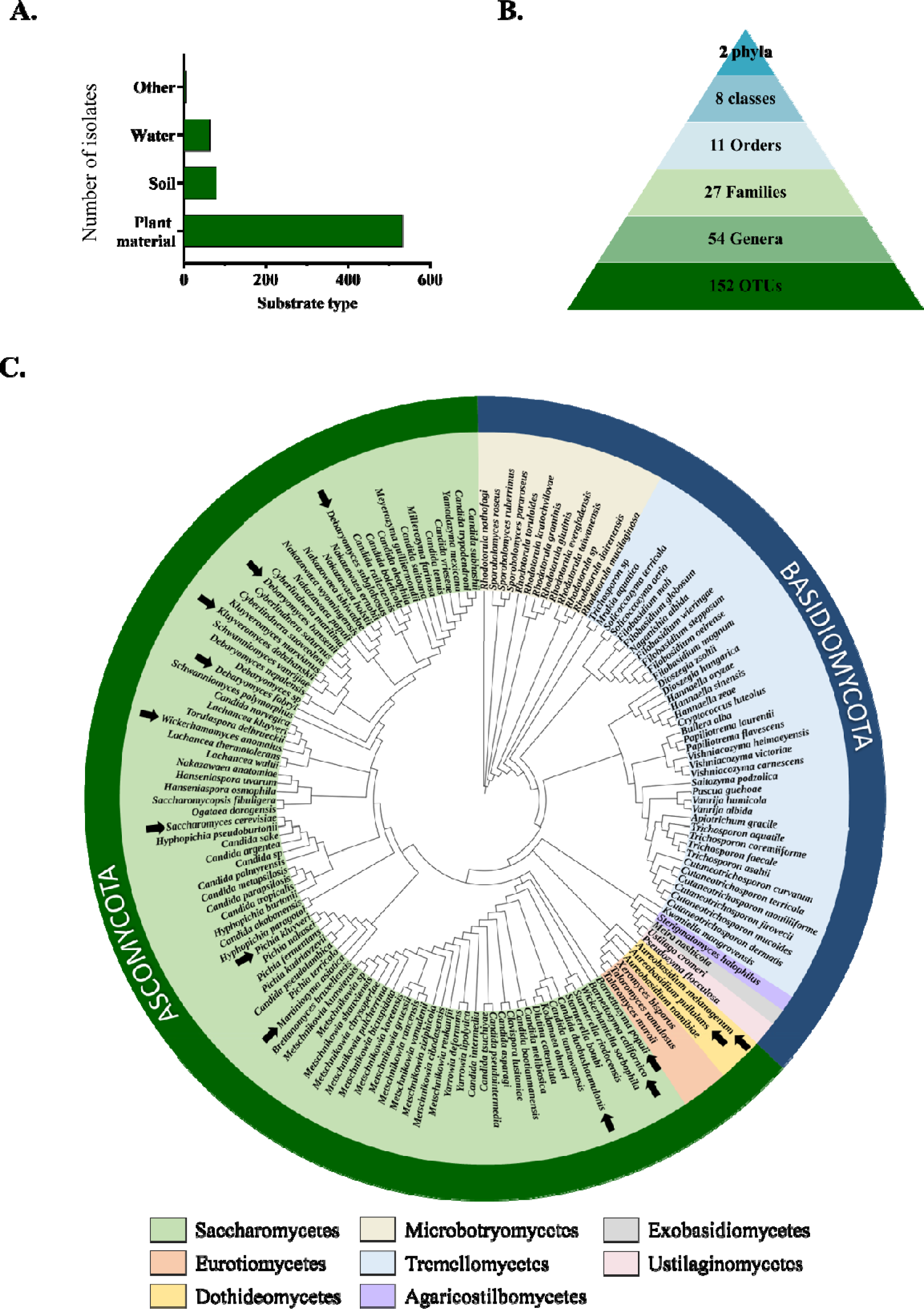
Overview on the species-diversity within the wild yeast collection. **A.** Number of yeast isolates from the collection classified per substrate type. **B.** Taxonomic diversity of the wild yeast collection. **C.** Phylogenetic tree built from multiple sequence alignment of ITS reads. The different phyla and families are highlighted. The arrows represent species that secreted a protein with antifungal properties.

Next we determined the species diversity within the yeast collection using the combined ITS1 and ITS2 amplicon (between 250 to 1,800 bp in lenght^69^) from the Internal Transcribed Spacers is often used to reveal species-identity and phylogenetic relationships^70^. Considering cost and workload, we applied a two-step approach, where we first used a pooled sequencing approach to identify which species were present in our collection, followed by individually sequencing isolates of interest later on.

For the first step, we pooled the collection from 96-well plates into four different samples; specifically two plates were pooled and sequenced together per sample, meaning that maximum 192 isolates were sequenced per 20,000 reads to increase cover and resolution by sample. After pooling, we extracted the DNA via regular ethanol precipitation and obtained DNA concentrations ranging from 25 – 40 µg/mL (**Figure 1C**). Samples were sent for PacBio sequencing, which allowed us to obtain long reads with sufficient depth do determine the bulk biodiversity within our collection. PacBio sequencing yielded a total of 1,199 Amplicon Sequence Variants (ASVs) **(Supplementary Table 4).** Each ASV was linked to the first hit with the highest coverage and similarity corresponding to a known yeast species using BLAST. ASVs ascribed to filamentous fungi, higher fungi or uncultured fungi were assumed to be contamination and therefore discarded.

Afterwards, ASVs ascribed to the same species were combined into a total of 150 operational taxonomic units (OTUs)^71^ (**Supplementary Table 5**). The reads length obtained from sequencing ranged from 104 to 956 bp (**Supplementary** Figure 1). Two additional OTUs – *Candida duobushaemulonis* and *Martiniozyma asiatica* – were later identified during individual Sanger sequencing of the D1/D2 domain and the ITS1 of killer yeast candidates and added to the list. The genomic DNA or the ITS amplicon of these species might have been below the detection limit in the PacBio sequencing sample.

In sum, we identified 152 different OTUs in our collection belonging to 54 different yeast genera distributed across 27 families, eleven orders, eight classes and two phyla (**Figure 2B** and **Supplementary Table 5**): Ascomycota (63%) and Basidiomycota (37%). As expected, most yeasts belonged to the order Saccharomycetales (59%); although other taxonomical categories were present as well. The collection contained dimorphic genera such as *Aureobasidium*^72^*, Trichosporon*)^73^ and *Sporobolomyces*^73^. Similar to previous yeast prospecting efforts, our collection contained species known as opportunistic human pathogens such as *Candida parapsilosis, Aureobasidium melanogenum* and *Pichia kudriavzevii* supporting that these yeasts are also associated with the natural environment^74,75^.

One representative ITS sequence from each OTU – chosen haphazardly – were used to create an alignment and build a phylogenetic tree (**Figure 2C).** In the case of *M. asiatica* and *C. duobushaemulonis*, the full ITS sequence of representative strain bank isolates *M. asiatica* CBS 10863 and *C. duobushaemulonis* CBS 7798 were used to increase the overlapping regions and allow for higher sequence homology and hence proper phylogeny.

### 31% of the yeast isolates show antifungal activity with a variety of target spectra

Next we screened the yeast collection for antagonistic activity against our panel of eight target yeast species. *S. cerevisiae* NCYC232 (natural producer of K1 killer toxin^76^) and *Williopsis mrakii* NCYC2251 (natural producer of HM-1 killer toxin^47^) were used as positive controls. Of note, K1 killer toxin did not show antifungal activity against every sensitive strain used, and was therefore excluded from further screening assays. The non-antagonistic *S. cerevisiae* laboratory strain BY4714 was included as a negative control. Every plate was replicated (in triplicates) on a screening plate containing an agar-embedded target yeast and methylene blue was added to the agar for a better readout^60^. Activity was judged by a growth inhibition halo around the isolate (**Figure 3A**) to better visualize the presence of a growth inhibition halo. As most yeast killer toxins showed optimal fungicidal activity at acidic pH and ambient temperature^77–82^, we performed this first screening assay at pH 4.0 and 25 °C.

**Figure 3.**
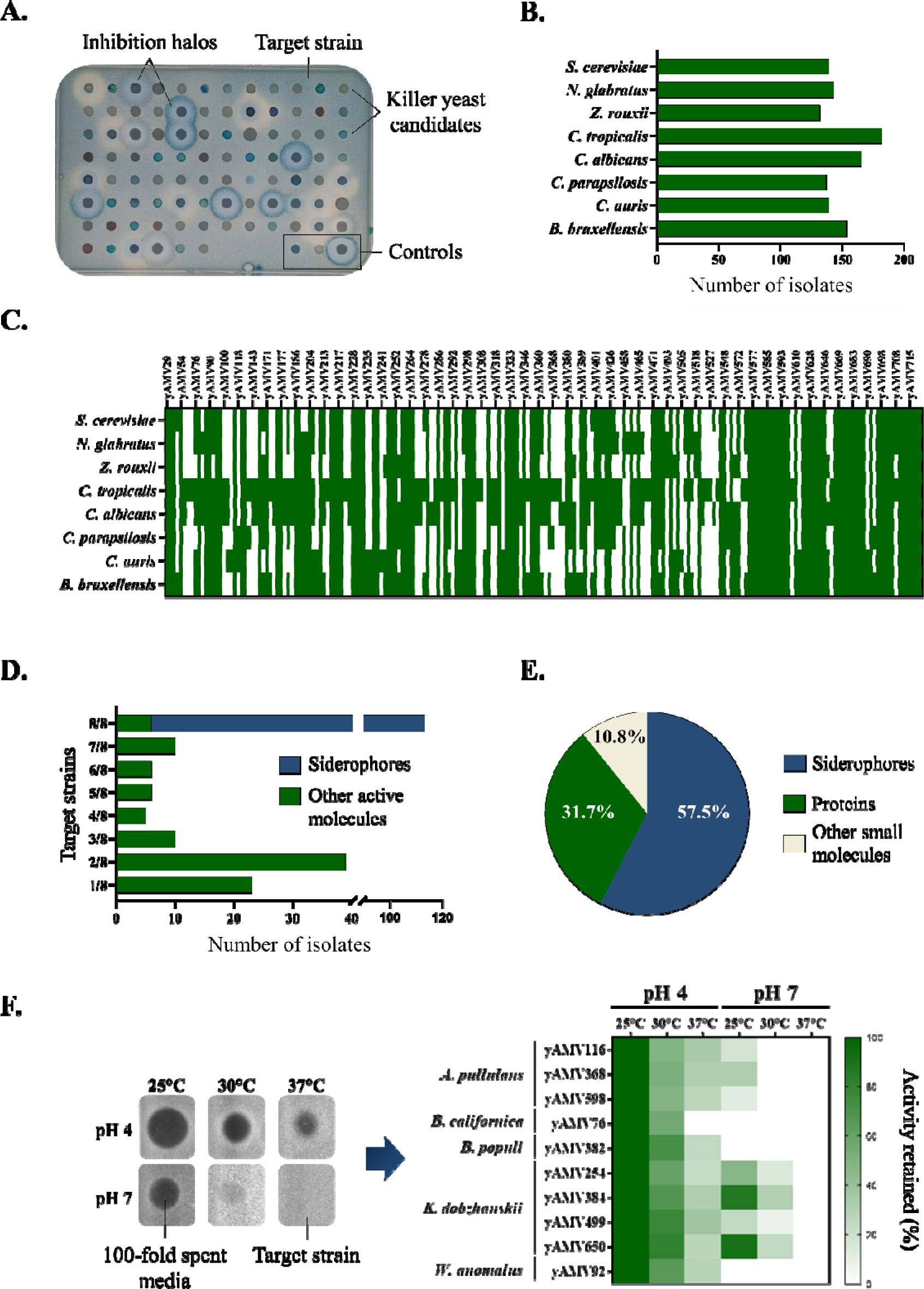
Overview on antifungal capacity within the wild yeast collection. **A.** Screening assay for antifungal activity. The target strain was embedded in the agar and the killer yeast candidates were spotted on top using a manual pinner. The yeast collection was divided into eight source plates screened against eight target strains used in this work. The controls were, from left to right, *S. cerevisiae* BY4741 (negative control), *S. cerevisiae* NCYC232 (K1 toxin producer) and *W. mrakii* NCYC2251 (HM-1 toxin producer). **B.** Number of isolates displaying activity against each target strain. **C.** Antifungal activity of every isolate against the eight target strains. The assay was performed in triplicate and the heatmaps depicts representative yes/no results. **D.** Overview on the various target spectra. Each column depicts how many yeasts were active against a certain number of target strains. For example, >20 yeasts were active against only one out of the eight yeasts of our target panel and >100 yeasts were active against all eight out of the eight yeasts and are thus broad spectrum. The colour code distinguishes between siderophore producers and “other active molecules”, where all siderophore producers were active against all eight target strains. **E.** Distribution of antifungal isolates classified by class of compound produced. The percentage shown is calculated based on active yeasts. **F.** Test for activity of pre-selected isolates under different application-relevant pH and temperature conditions using 100 – fold spent media. Measurements of halo sizes were normalized to values obtained at pH 4 and 25 °C (set to 100%), the reported values are the mean of triplicates and the standard deviation is reported in **Supplementary table 8**.

212 out of the 681 yeast isolates (31%) displayed an antagonistic effect against at least one of the eight targets (**Figure 3B**, **Figure 3C** and **Supplementary Table 1)**. Specifically, 132 isolates (62.3% of the active yeast) inhibited the growth of *Z. rouxii;* 137 isolates (64.6%) inhibited the growth of *C. parapsilosis;* 139 isolates inhibited the growth of *S. cerevisiae*, 139 isolates the growth of *C. auris* (65.6%); 143 isolates (67.4%) inhibited the growth of *N. glabratus*; 154 isolates (72.6%) the growth of *B. bruxellensis*; 165 isolates (77.8%) the growth of *C. albicans* and 182 inhibited the growth of *C. tropicalis* (85.8%). In sum, our collection yielded hundreds of yeast with activity against the target strains from our panel with *C. tropicalis* and *C. albicans* being overall the most susceptible targets (**Figures 3B and 3C**).

From those 212 active yeasts, and within the detection limit of our high-throughput assay, we observed a diverse set of target ranges **(Figure 3D)**. 113 isolates (53%) showed broad-spectrum activity inhibiting all targets tested; 76 isolates (36%) showed some degree of specificity, where 39 (18%) targeted two out of eight strains, ten (5%) targeted three strains, five (2%) targeted four strains, six (3%) targeted five strains, other six (3%) targeted six strains and ten (5%) targeted seven strains; finally, 23 isolates (11%) showed narrow- spectrum species-specific activity, inhibiting only one target yeasts of our panel.

Overall, *C. tropicalis* and *C. albicans* were not only the most targeted species, but several isolates showed exclusive activity only against those two species (for example, yAMV173 and yAMV178 – see **Supplementary Table 1**). 70% of species-specific isolates (16 isolates in total) were active exclusively against *C. tropicalis*.

Overall, the screen showed that yeasts with a variety of targeting profiles can be found in the environment.

### 107 isolates (half the active yeasts) show iron-dependent antagonism via the production of pulcherriminic acid-like iron siderophores

The 212 isolates that had shown a growth inhibition halo were re-arrayed into three 96-well plates and subjected to the next phase of screening, which aimed at identifying the type of molecule class and/or antagonistic mechanism involved (**Figure 1B**).

Yeasts naturally produce different compounds with antifungal activity^38–40^, one of them being iron-chelating siderophores. The most well-studied yeast-based iron siderophores are rhodotorulic acid, produced by *Rhodotorula* species (Basidiomycota)^83^ and pulcherriminc acid, produced by *Metschnikowia* and *Kluyveromyces* species (Ascomycota)^84–86^. While the presumed function of rhodotorulic acid production is scavenging of iron, with its production being repressed under high concentrations of iron^87,88^; production of pulcherriminic acid is discussed to rather serve for iron monopolisation^86^, *i. e.,* increasing concentrations of iron resulted in higher concentrations of the intracellular iron-pulcherriminic acid complex^89–91^ with a darker maroon-coloured cell pigmentation of the producer^92,93^.

We developed an assay that would rapidly allow to categorize those yeasts that antagonize fungi via production of iron siderophores and discern within rhodotorulic acid producers and pulcherriminic like producers based on the colour phenotype. The assay was based on pinning the isolates on plates with trace amounts of iron (control plate) and with iron supplemented (20 µg/mL FeCl_3_), the idea being that the growth inhibition halo of an iron-siderophore producing yeast would disappear in the presence of additional iron, while the growth inhibition halo due to a compound with a different mechanism of action would remain (**Supplementary** Figure 2). As controls, we used *S.* cerevisiae BY4741 (no antagonism), HM-1 killer toxin natural producer (iron-independent antagonism) and our isolate yAMV737, identified as *M. pulcherrima* via Sanger sequencing of the ITS1 region. This assay was performed in triplicates.

Using this screen, 107 of the 212 isolates were found to antagonise via iron monopolization, showing no inhibition halo with FeCl_3_ supplementation and a darker maroon-like cellular pigmentation, which indicated the production of pulcherriminic acid-like iron siderophores.

Although not all broad-spectrum isolates turned out to produce iron siderophores, all siderophore-producing yeasts from the collection showed broad-spectrum activity, targeting all yeasts tested (**Figure 3D**). This result is consistent with the hypothesis that these yeasts antagonize via depleting an essential nutrient – in this case iron – from the media, thus inhibiting in a species-independent manner. For future bioprospecting efforts focussed on one class of molecule, the high throughput screening and iron depletion assay could be combined. From the remaining yeasts, 79 kept the antifungal activity with additional FeCl_3_ and 26 yeasts, that had displayed antagonism in the first screen, stopped showing a growth inhibition halo (**Supplementary Table 6**).

### The presence of iron during enrichment has an effect on the isolation of yeast-based antifungal compounds

When consolidating the yeast collection, we started screening for activity before finishing the sampling. When screening the first batch of yeasts (yAMV1 – yAMV370), we observed that 45% of the active yeasts (15% of all yeasts), were iron-monopolizers. As for the purpose of this study, we aimed at finding as many true killer yeasts as possible, we supplemented the enrichment media with 20 µg/mL FeCl_3_ for the consolidation of the second part of the collection (yAMV371 – yAMV729, **Supplementary Table 1**). We reasoned that the iron supplementation would avoid competition during the enrichment and thus prevent out-competition of killer yeasts. However, 16% of all yeasts isolated were siderophore producers, similarly to the results obtained previously. Surprisingly, when looking at the percentage of iron-dependent antagonistic yeasts within the pool of active yeast isolates, the percentage of siderophore producers was 56%, even higher when compared to the 45% observed before (**Supplementary** Figure 3). Counterintuitively, iron supplementation of the enrichment media had no effect on isolating iron-monopolizing yeasts but was counterproductive when aiming to isolate yeasts producing other antifungal molecules. Though this finding is not statistically significant (Χ^2^(1, *N* = 200) = 1.0026, *p =* .32), it is an observation worth considering for future bioprospecting studies.

### 59 yeast isolates produce a high molecular weight compound with antifungal activity

In the next step, the 79 yeast isolates that showed no iron-dependent phenotype were tested for the secretion of a higher-molecular weight compound (>3 kDa) – likely proteins – by performing a size-exclusion assay on their spent media.

The assay started with culturing the yeasts in 5 mL culture volume for 24 hours, removing the cells via centrifugation, followed by concentrating their sterile-filtered spent media 10 – 40 fold with centrifugal filter units containing a membrane with 3 KDa size cut-off. This way, only compounds with a molecular mass larger than approximately 3 KDa were retained in the concentrate. After this, the concentrated spent media was assayed for inhibitory activity by a spot assay on 150 mm round Petri dishes (**Supplementary** Figure 4). From these 79 isolates, 59 isolates produced a compound larger than 3 KDa responsible for the activity (**Figure 3E**). That meant 8.7% of yeast isolates from the collection (31.7 % of the active ones) secreted a proteinaceous compound with antifungal activity and were thus candidates to be killer toxin producers.

### The 59 killer yeast candidates cluster into ten different OTUs

In order to identify the 59 killer yeast candidates individually at species level, we amplified and Sanger sequenced their ITS1 region, or their D1/D2 domain when the ITS1 was not successfully amplified. Sequences were blasted and defined as the species from the first hit with highest query coverage (%) and sequence similarity (%) (**Supplementary Table 7)**.

For 15 isolates, the 97% similarity threshold to the closest related ITS was not reached. We grouped them into the species corresponding to the highest similarity match. Moreover, some sequences retrieved more than one species with the same percentage of sequence similarity and query coverage when blasted. Here, we grouped them into the species with the most entries. Specifically, isolates that yielded the same coverage for *Aureobasidium melanogenum* and *Aureobasidium pullulans* were referred to as *Aureobasidium pullulans* and all species belonging to the genus *Debaryomyces* were classified as *D. hansenii,* although the three species *Debaryomyces hansenii, Debaryomyces fabry* and *Debaryomyces subglobosus* were retrieved from our search^94^.

The killer yeast candidates eventually grouped into ten species with 32 isolates of *Debaryomcyes hansenii*, seven isolates of *Wickerhamomyces anomalus,* seven isolates of *Barnettozyma californica,* four isolates of *Kluyveromyces dobzhanskii,*, three isolates of *Aureobasidium pullulans,* two isolates of *Pichia kluyveri* one isolate of *Martiniozyma asiatica,* one isolate of *Candida duobushaemulonis,* one isolate of *Saccharomyces cerevisiae* and one isolate of *Barnettozyma populi*. The query coverage, sequence similarity and species ascribed to each isolate are available in **Supplementary Table 7**. Apart from *A. pullulans* (Class: *Dothideomycetes*, Order: *Dothideales*), all killer yeast candidates belonged to the class *Saccharomycetes* order *Saccharomycetales* (**Supplementary Table 5**). *A. pullulans*^68^*, B. californica*^95^*, D. hansenii*^96^*, K. dobzhanskii*^97^*, P. kluyveri*^98^*, S. cerevisiae*^76^ and *W. anomalus*^79^ isolates had previously been reported as killer yeasts. To the best of our knowledge, it is the first time that *B. populi, M. asiatica* and *C. duobushaemulonis* have been reported to produce a protein with antifungal properties.

### Some killer yeast candidates retain their activity at neutral pH and 30 ***°C***

Most yeast killer toxins report an optimal killing activity at pH 4 – 4.5 and 20 – 25 °C^77–82^. We tested if the compounds produced by our killer yeast candidates were active at different temperatures and pH values. We started the assessment using a high-throughput halo assay similar to the one used for the initial screen of the full collection with the media adjusted to different pH values (pH 4.0 and pH 7.0) and incubated at different temperatures (25 °C, 30 °C and 37 °C). However, many yeast isolates showed growth defects at higher temperature and / or pH, compromising the reproducibility of the results. Therefore we used this assay as a pre-screen to identify promising candidates that retained activity across temperature and pH conditions. According to the pre-screen, ten isolates were chosen, specifically four *K. dobzhanskii* isolates, three *A. pullulans* isolates, one *B. californica* isolate, one *B. populi* and one *W. anomalus* isolate. They were cultured this time in 100 mL; their spent media was 100 – fold concentrated and used in a spot assay (10 µL) for activity against *C. tropicalis* (**Figure 3F, Supplementary Table 8**).

For all 10 concentrates, the highest inhibitory activity was measured at pH 4 and 25 °C, so the measurements of all conditions were normalized to the halos obtained at these conditions. Some isolates retained some activity at 30°C and, interestingly, all four proteins produced by *K. dobzhanskii* (yAMV254, yAMV384, yAMV499 and yAMV650) retained some activity at pH 7, specially yAMV384 and yAMV650 (**Figure 3F)**. Activity at neutral pH had also been found previously for zymocin, a well-studied yeast killer toxin produced by *Kluyveromyces lactis*^48^.

### The killer yeast candidates cluster in 46 different same-species activity profiles

It is known that different strains from the same species can secrete different killer toxins (*e.g.* K1, K2 and K28 from *S. cerevisiae*^99–101^ or KP4 and KP6 from *Ustilago maydis*^102,103^). The fact that all same-species killer yeast candidates were isolated from different substrates, at different dates and distinct locations gave us confidence that all samples were independent isolates and not clones from the same environmental sample or contaminations that occurred during the screening. Nevertheless, we wanted to compare the inhibitory profile of all same-species isolates and cluster them into groups of distinct patterns, potentially indicating the production of different proteins responsible for the antifungal activity. To refine this comparison, we quantified the halo area size – as larger halos could indicate a more effective molecule being secreted or simply more of the same molecule – and compared the ITS1 sequences obtained for species identification. This time we performed a similar halo assay as before but spaced the yeasts across the 96-well format to be able to get distinct non-overlapping measurable halos (**Supplementary** Figure 5A), a concern specially when screening against *B. bruxellensis* which yielded very large halos most likely due to its slow growth. The yeast yAMV388 stopped producing a protein with antifungal activity in this phase of the screening. The three *A. pullulans* isolates were screened individually via spotting 5 µL of an OD_600_ normalized cell-suspension to 50 units, as they showed expansive growth and did not grow reliably when pinned with the manual 96-array pinner (**Supplementary** Figure 5B). **Supplementary Tables 7 and 9** show the measurement values of halo area size for all yeasts, the latter containing the values for *A. pullulans* yeasts.

Eventually the killer yeast candidates grouped into 46 different same-species-activity profiles based on halo size, target range and ITS1 sequence (**Figure 4)**.

**Figure 4.**
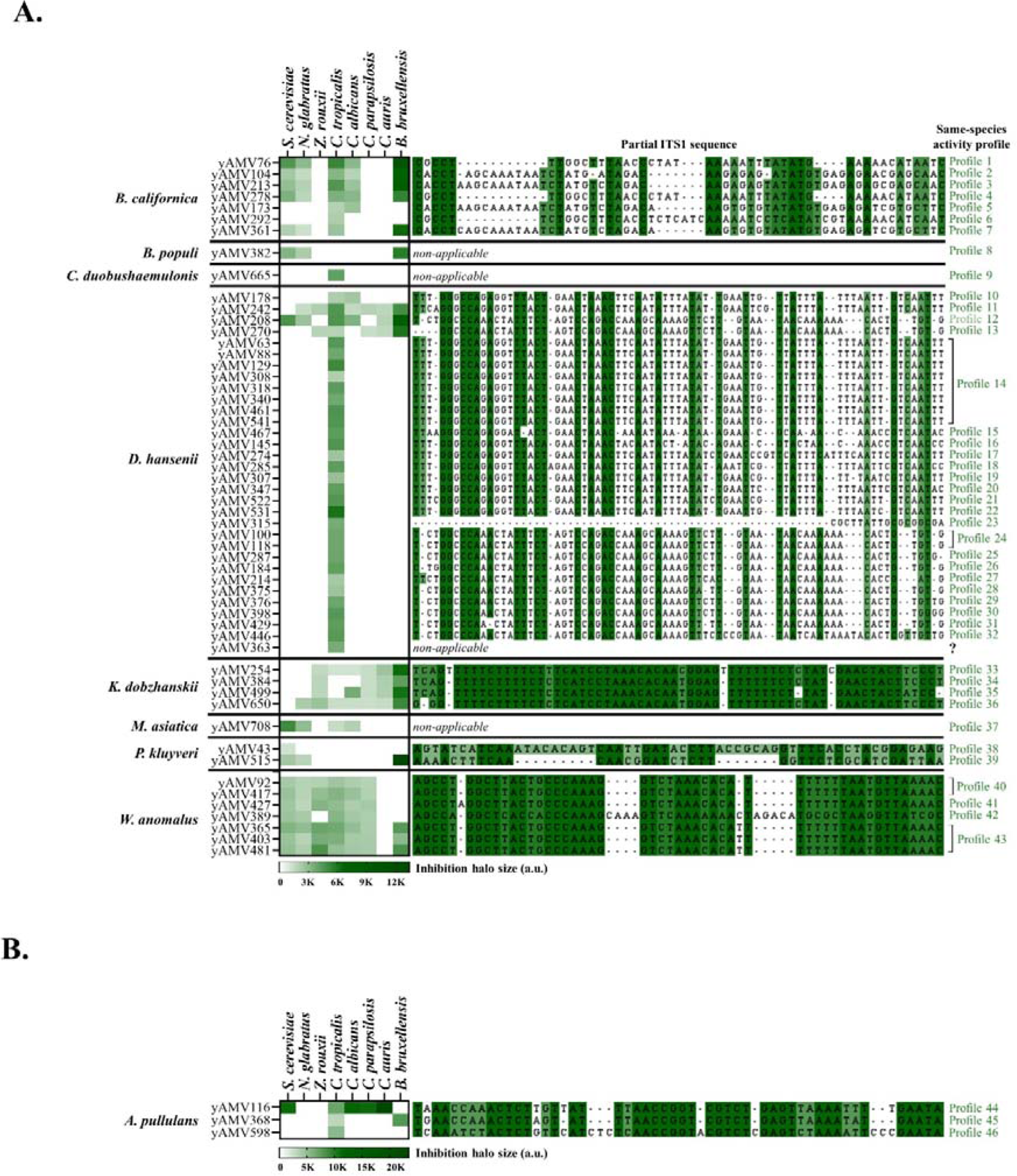
Activity profiles of the yeast isolates. On the left, the halo size measurements of each yeast isolate grouped by species is given. On the right, a multiple sequence alignment of the partial sequence of the ITS1 regions of same-species isolates is given. Based on the halo size, activity profile and ITS1 sequence, at least 43 different species-related activity profiles were determined. Sequences of isolates where no other same-species isolate was present and isolates whose species-level were identified by D1/D2 sequence are shown as *not applicable*. The question mark (?) stands for an isolate identified as *D. hansenii* by D1/D2 sequencing that, consequently, cannot be compared to other *D. hansenii* isolates. **A.** Results of yeast isolates tested using the manual pinner. B. Results of *A. pullulans* isolates, screened separately due to their expansive growth. The heatmaps depict the mean of triplicate experiments and the standard deviation is given in **Supplementary tables 7 (A)** and **9 (B)**.

For *B. californica,* yAMV76, yAMV104, yAMV213 and yAMV278 presented the same activity profile (**Figure 4A**). However, yAMV104 and yAMV213 had distinct ITS1 sequences. Moreover, yAMV76 displayed remarkably larger halos against *B. bruxellensis* in comparison to yAMV278. All isolates constituted seven individual profiles.

For *D. hansenii,* several killer yeast candidates showed a very similar antifungal phenotype and identical ITS1 sequences (**Figure 4A**), pointing to closely related isolates possibly producing the same compound. Nevertheless, the 32 isolates could be grouped in 23 profiles based on different ITS1 sequences. Interestingly, most isolates were highly specific against *C. tropicalis.* In the case of yAMV363, we could not amplify the ITS1 but rather the D1/D2 domain; consequently, that yeast could not be compared to the other same-species isolates and it remains unclear whether it might compound another activity profile.

For *W. anomalus,* four different activity profiles were identified based on target range and ITS sequences. Isolates yAMV92 and yAMV417 and isolates yAMV365, yAMV403 and yAMV481, were grouped based together on these characteristics.

For *K. dobzhanskii* and *P. kluyveri,* all isolates showed unique target range and ITS sequence, hinting to the production of different antifungal compound.

For *B. populi, C. duobushahemulonis* and *M. asiatica*, only one isolate per species was identified in the collection, so no comparison with ITS sequences was necessary, all isolates constituted different individual activity profiles (**Figure 4A**).

Finally for *A. pullulans* isolates, assays were performed separately as these isolates showed expansive growth and not allowing for precise halo-size measurements (**Supplementary** Figure 5B). All three yeasts showed unique inhibitory phenotypes and sequences, constituting three different same-species activity profiles (**Figure 4B**).

It should be noted that our results using the manual pinner yielded an ‘apparent’ target spectrum based on the detection limit of the assay. For *A. pullulans* we observed a more refined target spectrum when concentrating the spent culture media, meaning that inhibitory activity could be observed against targets that would not be visibly inhibited in the pinning assay (**Supplementary** Figures 5B and 6).

We also compared isolates with a similar inhibitory profile belonging to different species. Yeasts with a similar target specificity might produce proteins that share binding receptors, targets, or overall mechanisms of action. We grouped the isolates showing a similar inhibitory profile in four different species-independent activity profiles (**Supplementary** Figure 7). In short, yAMV173 (*B. californica*) and yAMV178 (*D. hansenii*) both showed inhibitory activity against *C. tropicalis* and *C. albicans* (Profile 1); yAMV242 (*D. hansenii*) and yAMV650 (*K. dobhanskii*) antagonised every target strain of our panel except for *S. cerevisiae* (Profile 2); yAMV382 (*B. populi*) and yAMV515 (*P. kluyveri*) were active against *S. cerevisiae, N. glabratus* and *B. bruxellensis* (Profile 3); and, as mentioned above, multiple *D. hansenii* isolates targeted *C. tropicalis* specifically, a common characteristic with yAMV292 (*B. californica*) and yAMV665 (*C. duobushaemulonis*) (Profile 4).

### Several killer yeast candidates inhibit mycelial growth or conidiation of B. cinerea

Next we tested the 59 killer yeast candidates against the filamentous plant pathogen *B. cinerea*. We placed pre-grown mycelial plugs of the fungus in the centre of an agar plate, followed by spotting 10 µL of a cell suspension of the killer yeast candidates three times on the plate surrounding the fungal plug (**Figure 5A**). After 7 days of incubation, three different phenotypes were observed (**Figure 5B** and **Supplementary Table 7**): no inhibition, inhibition of conidiation and inhibition of mycelial growth. 42 isolates did not cause any inhibition on the plant pathogen. 14 isolates showed absence of conidiophore formation on plate but mycelial growth could be observed around yeast cells. Finally, only three isolates inhibited completely the mycelial growth of *B. cinerea* **(Figure 5B)**. The inhibition of conidiation phenotype was scattered across species, while the growth inhibition phenotype was only observed in the three *A. pullulans* killer yeast candidates (**Figure 5C**). Some species showed only one phenotype (*e. g*. *S. cerevisiae* and *M. asiatica*) but, as the collection contained only one active isolate for those species, no conclusions could be drawn from this.

**Figure 5.**
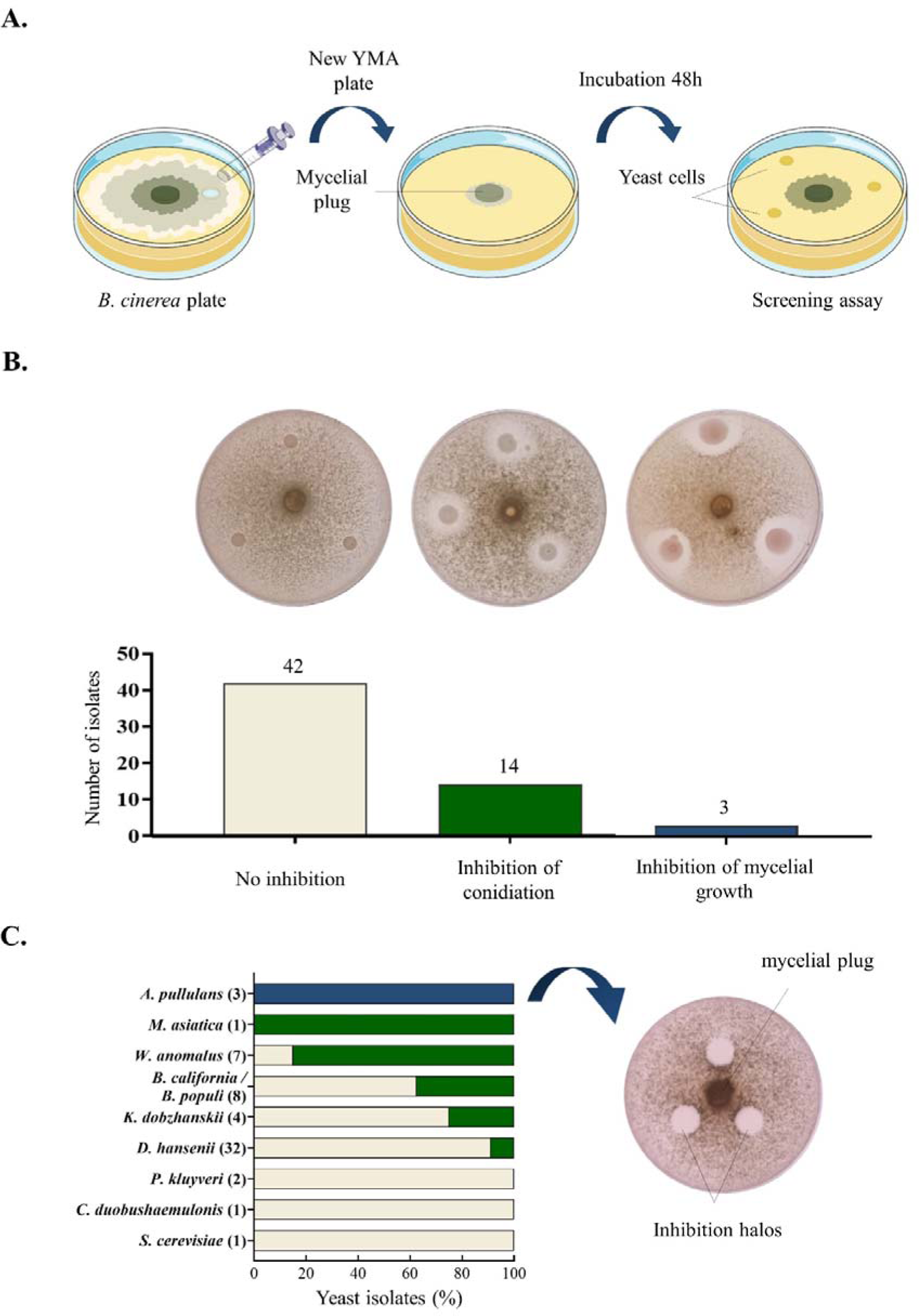
Antagonistic effect on the plant pathogen *B. cinerea*. **A.** Workflow of the screening assay. **B.** Number of yeast isolates per phenotype observed (from left to right): no inhibition, inhibition of conidiation and inhibition of mycelial growth. Representative picture on top of each bar graph **C.** Distribution of inhibitory phenotype per species. Number of isolates per species between brackets. On the right, 200 – fold supernatant of *A. pullulans* isolates tested for mycelial growth inhibition.

To verify that also this anti – *B. cinera* activity of the three *A. pullulans* isolates was due to a high-molecular weight compound, we tested their 200 – fold concentrated spent culture media against *B. cinerea* by spotting 5 µL similarly to how the screening was performed with the corresponding cells. All three concentrates, showed the same inhibitory activity that had been observed when testing the cells (**Figure 5C**).

### Yeast producing antifungal compounds are present in daily consumed products

Several active yeasts from our collection were GRAS organisms and displayed antifungal activity against diverse pathogens under application-relevant conditions. For future product development in food and biocontrol, GRAS levels and food-safety of the actual compounds secreted by these yeast would be a first cause for concern. In order to test the prevalence of yeast-based antifungal compounds in our food chain, we tested how many product-associated yeast isolates would show an antifungal phenotype. The prescence of these compunds would support the notion that humans are already consuming yeast-based antifungal compounds. Thus, we consolidated a set of 113 yeasts from daily-use commercial products, using the same isolation procedure as before. These products included fruits from different markets (81 isolates), yogurt (1 isolate), kombucha (2 isolates) and commercial wheat beers (9 isolates), as well as commercially available packaged baking yeast (10 vendors) and brewing yeast (10 vendors) (**Figure 6A** and **Supplementary Table 10**). We screened this collection for action against a subset of our panel of target yeast, specifically *S. cerevisiae, N. glabratus, C. tropicalis* and *B. bruxellensis.* A total of 26 yeasts (23% of the collection) displayed antifungal activity against at least one target strain (**Supplementary Table 10**). Interestingly, 30% of the tested brewing yeasts and 78% of yeast isolated from commercial wheat beers showed an antifungal phenotype against at least one species **(Figure 6B)**. Specifically, 16 isolates were active against *S. cerevisiae*, 14 against *C. tropicalis*, 13 against *B. bruxellensis*, and 11 against *N. glabratus* **(Figure 6C)**. In sum, our screen supports the hypothesis that antagonistic yeasts are ubiquitous, even in the food chain. While in-depth molecular characterization of the secreted compounds would be necessary to determine GRAS levels and food-safety, the discovery of antifungal yeasts in daily used products hints towards the fact that humans might already consume yeast-based antifungal compounds.

**Figure 6.**
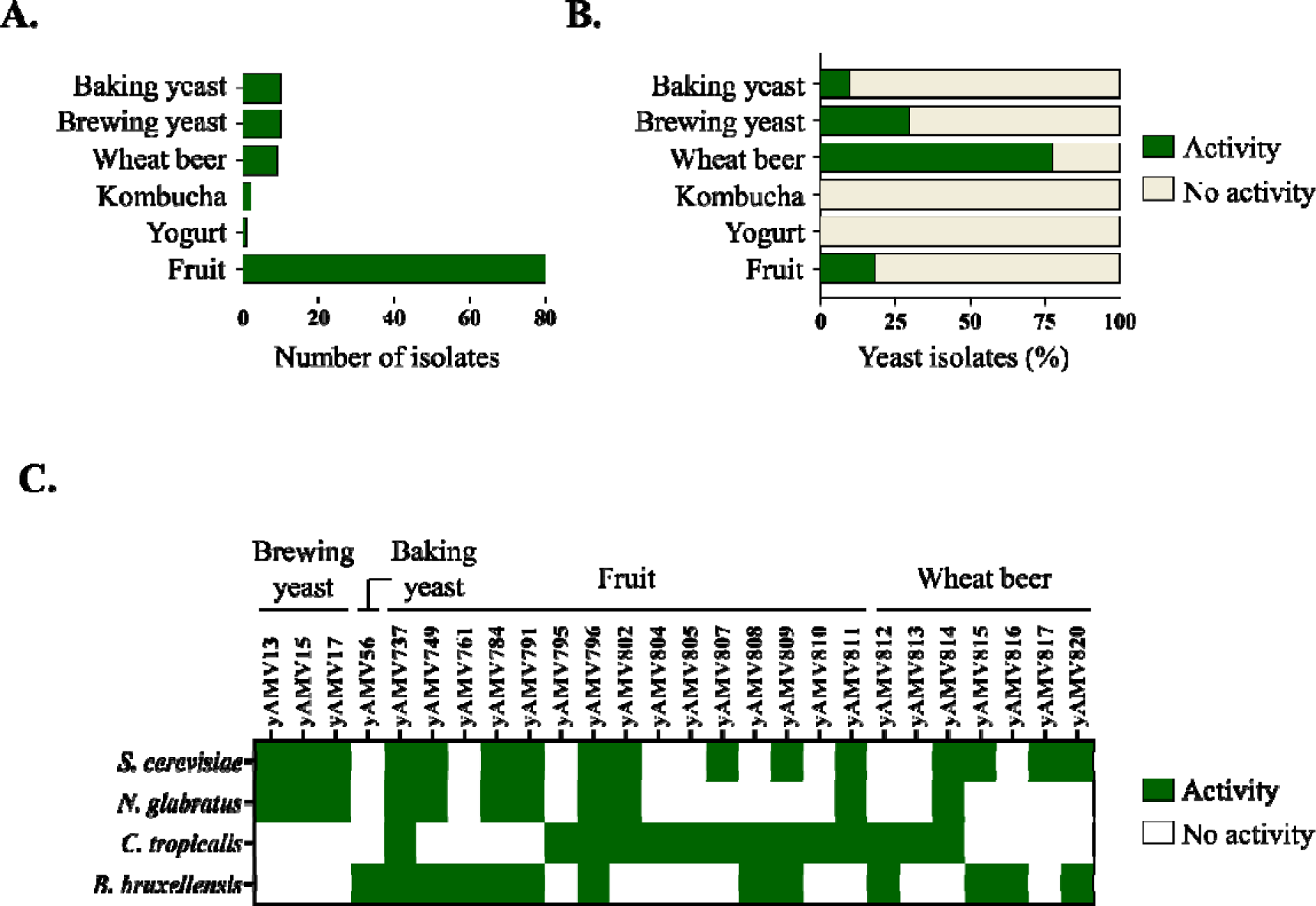
Screening of antifungal activity in daily consumed products. **A.** Number of isolates screened per type of product. **B.** Percentage of isolates with antifungal activity per type of product. **C.** Activity profile for all active yeasts tested against four different target strains. The assay was performed in triplicate and the heatmaps depict representative yes/no results.

### Calibration of antifungal activity with a commercial calibrant

Until now we had reported all halo sizes in arbitrary units. Due to the many factors that contribute to the size of a growth inhibition halo, we wanted to calibrate the halo size to an internal standard that could be used by any researcher, allowing to compare results across laboratories and studies. In addition, we wanted to create a context for the performance of these yeast compounds in comparison to a widely used antifungal. After testing a few antifungals (data not shown), we decided on the commercially available antifungal micafungin. It showed well-defined halos in spot assays, acted against the entire panel of eight target yeast and maintained activity across a wide range of pH and temperature values, although it showed highest activity at neutral pH.

For calibration, we spotted increasing concentrations of micafungin next to the target yeast or their concentrated spent media; our calibration curve consisted of 5 µL of 0.5, 1, 2, 4 and 8 µg/mL (see **Supplementary** Figure 5). We chose these values based on pre-testing various concentrations (data not shown) and selecting the ones that would yield a measurable range of halos within our assay and based on reported MIC values; for example, for multiple *Candida* spp, MIC is generally lower than 2 µg/mL.^104–106^ Plotting the halo sizes over concentration yielded a calibration curve that allowed to translate the yeast-based inhibition halos into equivalent micafungin units. We calibrated two assays, a spot assay with live yeast cells and a spot assay with 200 – fold concentrated spent media. The latter was performed only for the three *A. pullulans* isolates yAMV116, yAMV368 and yAMV598, as those isolates did not form nice round spots but rather showed expansive growth. Of note, it was important for this assay to run a calibration curve on each screening plate as the size of the halo was dependent on the target strain, the exact inoculum (OD_600_ needs to be accurately normalized), the pH and temperature. The halo size of each killer yeast candidate in equivalent micafungin units is depicted in **Supplementary** Figures 6 and 8 and the values are listed in **Supplementary Tables 9 and 11**.

Every killer yeast active against *C. parapsilosis* showed a halo that, although small when analysing absolute values, was equivalent to ≥2 µg/mL micafungin (**Supplementary** Figure 8). For other *Candida* species, our yeast targeted within 0.5 – 2 µg/mL micafungin equivalents, which is within the range used as treatment for susceptible *Candida* species^104^.

The isolates yAMV116, yAMV368 and yAMV598, from *A. pullulans,* all three reported an activity equivalent to 5 – 30 µg/mL micafungin when their 200 – fold spent media was tested (**Supplementary** Figure 6). Halos produced by cells were measured but their comparison to micafungin was not taken into consideration due to their expansive growth.

## Discussion

### Assessment of the workflow

Here we present a systematic workflow to isolate antifungal yeasts from the environment, with a specific focus on isolating killer yeast candidates with application-relevant target spectra and temperature and pH performance metrics. This workflow can be adjusted to other class of yeast-based antifungal compounds according to the downstream applications. Our work led to the consolidation of a 681-membered yeast collection comprising 152 OTUs, from which 59 independent killer yeast candidates with diverse activity and target profiles were isolated, as well as 212 siderophore producers. Killer yeast candidates belonged to 10 species with three species not having been reported before to secrete killer toxins. 46 of the killer yeast candidates potentially secrete different proteins, based on their activity profiles and ITS1 sequences. Our workflow highlights that a large number of inhibition-halo producing yeasts are siderophore producers, but that a combination of two simple screens (iron depletion and size exclusion) could be employed to discriminate between those and potential killer yeast candidates.

While we limited ourselves to specific screening conditions, which miss antifungal yeasts that target pathogens not represented in our panel or that require specific conditions for activity (*e.g.* high salt concentration,^107,108^ a specific pH^77–82^ or temperature^79–82^ or a transcriptional activator^109^), our workflow can be adapted to other pathogens and screening conditions that a future researcher is interested in.

Specific growth or selection conditions required for continuous production of a killer toxin could explain why some of our killer yeast candidates lost the inhibitory activity along our study.

Our workflow could be adapted to aim for other yeast-based classes of antifungal molecules. For example, a high-throughput screening with iron would yield the identification of iron siderophore producers. The colony pigmentation would give information on whether those are pulcherrimin-like or rhotodorulic-like siderophores.

In the future, the workflow needs to be extended by genetic and molecular characterization screens to identify the compounds responsible for antifungal action.

### Biodiversity of the yeast collection

We identified 152 different OTUs present in our 681-membered collection (4-fold redundancy). This diversity is slightly higher than a similar study undertaken in the US where 244 OTUs where identified in 1962 isolates (8-fold redundancy).^110^

Not every ASV sequence obtained from the pooled sequencing could be assigned to a known OTU based on the >97% sequence similarity threshold but individual sequencing of all yeasts would be necessary to verify the presence of novel species.

### Number of active yeasts and killer yeast candidates

The search for novel antifungal compounds from environmental yeasts has been made in numerous occasions.^50–52,55^ In our study, we found that 31% of our yeasts showed an antagonistic phenotype. This percentage is in line with prior findings in Brazil^51^ and Poland^55^, where 33% and 39% of naturally isolated yeasts showed fungal growth inhibition. However, different screening conditions and target strains were employed.

With respect to the occurrence of the killer yeast phenotype, 8.7% of our isolates turned killer yeast candidates. Although this result is in line with the 5 – 30% killer yeast frequency reported by other studies^111^ those relied on a simple halo assay as a readout,^51–57^ with no discrimination between siderophore producers, true killer yeast candidates and other smaller compounds. As such the frequency of true killer yeast is likely at the lower end of the 5-30% range.

### Applications in beverage preservation and biocontrol

The killer yeast candidates isolated in this study showed highest activity when tested at pH 4.0 and 25 °C. Those conditions are suitable for their use as biocontrol agents in agriculture or preservatives in the food and fermented drinks industry, either the yeast themselves or their produces compounds.

*B. bruxellensis* is a prevalent microbial contaminant in fermented beverages because of its slow growth and resilience at harsh conditions.^112–114^ Interestingly, several of our yeast isolates that targeted *B. bruxellensis* were inactive against *S. cerevisiae* within our assay’s detection limits (yAMV242, yAMV254, yAMV384, yAMV499 and yAMV650), yielding the potential for species-specific biocontrol of *B. bruxellensis* in beverages that are fermented by *S. cerevisiae,* as suggested before by others.^78^

In terms of biocontrol, our target pathogen *B. cinerea* (gray mold) is a major concern for agriculture and post-harvest control. Its primary way of transmission is by airborne asexual spores (conidia)^115^. Consequently, yeasts that inhibit the conidiophores formation present potential for bio-containment of infected plants. The selective inhibition of conidiation with no effect on mycelial growth has been previously observed^116^ and a recent list of key genes essential for conidiation but not for hyphal growth could serve as a starting point for further research.^117^

*A. pullulans* species have been previously reported to antagonize *B. cinerea* effectively^118–120^ where the inhibition was due to their chitinolytic activity.^68,121^ In fact, the company San Agrow© has released two products, Botector® and Blossom Protect®, whose formulations are based on *A. pullulans* isolates for the protection of fruits based on competition for nutrients and space. In the case of yAMV116, yAMV368 and yAMV598, the growth inhibition was caused at least partially by a secreted protein. Molecular and genetic studies would be necessary to determine if this activity is also due to a chitinase or a killer toxin.

### Medical applications

We isolated yeasts with a diverse set of targeting-specificities against the herein tested human fungal pathogens. Specifically *C. tropicalis* and *C. albicans* were very often targeted by our yeast isolates. As those yeasts are the most frequently isolated *Candida* species in candidiasis/candidemia patients in the Asia-Pacific-region,^122–124^ further research on novel antifungals with certain level of specificity towards these pathogenic yeasts could be promising for the development of fungal vaccines, drugs and other biopharmaceuticals.

While the micafungin equivalent values introduced in this study were aimed at calibration for inter-laboratory reproducibility, comparison of the activity of killer yeast with this antifungal could be used as an indicator for their efficacy. We note that our yeasts most likely have a different mode of action: micafungin inhibits fungal cell wall biosynthesis^125^ while yeast killer toxins present a wider range of killing mechanisms.^41–44^ Moreover, the conditions used for the different assays used an acidic pH for killer toxins;^77–82^ while the optimal pH of micafungin is between 6.2 and 6.9.^126^

Our killer yeast candidates targeted *Candida* species within 0.5 – 2 µg/mL micafungin equivalents, being ≥2 µg/mL micafungin equivalents in the case of *C. parapsilosis*. Those values are within the range used as treatment for susceptible *Candida* species.^104^ Micafungin and other echinocandins are the first choice for treatment treatment and prophylaxis against *C. parapsilosis*; ^127,128^ however, multiple resistant strains have been reported.^129,130^ Consequently, the use of yeast killer toxins could be a very promising alternative for biomedical treatments of candidemia and candidiasis patients with this fungal pathogen.

## Conclusion

Summarizing, our workflow delivered hundreds of yeast isolates with diverse target spectra and promising characteristics for applications in food, biocontrol and human health. Molecular and genetic characterization needs to follow to identify the underlying molecules and their biosynthesis.

Most of our isolates were siderophore producers, showing the importance of discriminating siderophore producers from true killer yeasts to estimate the real abundance of killer yeasts in the environment.

Focussing on killer yeast candidates, our study confirms that killer yeasts can be found across many yeast genera and environments. We show for the first time that many foods we consume every day already contain killer yeast. We show that killer yeast candidates exist with narrow and broad target spectra and profiles beyond low pH and low temperature.

In the future, our workflow and yeast collection could aid the identification of diverse yeast-based antifungals active against relevant pathogens.

Considerations and recommendations when using our workflow:

**1.** False negatives due to specific screening conditions: The workflow presented in this study yields yeasts that produce an active compound constitutively and which is active at our chosen screening conditions and against our target panel of pathogens. Different growth and screening conditions, active compounds or target species can be applied depending on the research objectives.
**2.** Limit of detection and “apparent” target range: The pinning assay we use to determine the target range of each antifungal yeast isolate has a specific limit of detection, based on the number of cells that get pinned, the susceptibility of the target strain and the potency of the secreted compound. Most toxins have been reported to be secreted in rather low amounts.^131–134^ Thus the reported target range is only the range as it appears from our assay. Although we show examples where concentrating the spent media and hence the active proteins helps in defining the target range, we argue that our assay is still suitable to report a certain specificity in targeting **(Supplementary** Figures 5B and 6).
**3.** Secretion of multiple compounds: Our workflow yields yeast-based secreted proteins that contribute at least partially to a fungal inhibitory phenotype. Some yeasts are known to secrete multiple compounds that contribute to their antagonism (*e.g. Metschnikowia* spp)^84^ but our workflow does not yet discriminate for that.
**4.** Panel of single-strain target yeasts: Here we only test a single strain per target species. Some yeasts show antifungal activity at a strain level.^135^ Follow up studies could test activity against a large panel of strains of a given species to determine the specificity at a strain level of an active yeast.
**5.** Influence of the media composition used for enrichment: We observed that the presence of supplemented iron did not increase the amount of siderophore-producing yeasts isolated but did decrease the amount of yeasts producing proteins and other active compounds. Although the results were not statistically significant, they are worth considering on future bioprospecting efforts.

## Contributions

AMV and SB designed the research. AMV performed the experiments. FT performed replicates for the pre-screen of killer yeast candidates at different pH and temperature. AMV drafted the manuscript. SB revised the manuscript. All authors have read and approved the final version of the manuscript.

## Acknowledgements

The authors would like to thank Vasiliki Tsousi and Marta Cardoso for their assistance during the collection and isolation of environmental samples, all friends and colleagues who donated environmental samples, Dr. Jan AKW Kiel for kindly donating yeasts isolated by students from the Rijksuniversiteit Groningen and Jie Xu from BMKGENE for his assistance during the ITS PacBio sequencing process.

