## Supplementary Tables 1-3; Supplementary Figures 1-8 for "The antifungal capacity of a 681-membered collection of environmental yeast isolates"

**Supplementary Table 1**. Yeast samples present in the collection with their location and material source. The fungal growth inhibition observed via halo assay is indicated for each background strain tested as 'yes' (active) and 'no' (not active).

**Supplementary Table 2.** Strains used in this study, characteristics, use and source.

| **Strain** | **Characteristics and use** | **Reference/source** |
| --- | --- | --- |
| *S. cerevisiae* BY4741 | BY4741 *MAT*a *leu2Δ0 met15Δ0 ura3Δ0 his3Δ1* – Target strain to test antifungal activity and negative control in screening assays | Brachmann *et al*. (1998)^43^ |
| *N. glabratus* ATCC2001 (formerly known as *C. glabrata*) | Human fungal pathogen.  Target strain to test antifungal activity | American Type Culture Collection |
| *C. tropicalis* NCYC4 | Human fungal pathogen.  Target strain to test antifungal activity | National Collection of Yeast Cultures |
| *C. parapsilosis*  NCYC601 | Human fungal pathogen.  Target strain to test antifungal activity | National Collection of Yeast Cultures |
| *C. auris* ATCC-MYA-5001 | Human fungal pathogen.  Target strain to test antifungal activity | American Type Culture Collection |
| *C. albicans*  NCYC597 | Human fungal pathogen.  Target strain to test antifungal activity | National Collection of Yeast Cultures |
| *Zygosaccharomyces rouxii* NCYC567 | Food and beverage spoiler.  Target strain to test antifungal activity | National Collection of Yeast Cultures |
| *Brettanomyces bruxellensis* NCYC370 | Beverage spoiler.  Target strain to test antifungal activity | National Collection of Yeast Cultures |
| *B. cinerea* CBS 125.27 | Plant pathogen. Target strain to test mycelial growth inhibition | Westerdijk Fungal Biodiversity Institute |
| *Williopsis mrakii* NCYC2251 | Natural producer of HM-1 killer toxin – positive control for antifungal activity assays | National Collection of Yeast Cultures |
| *S. cerevisiae* NCYC232 | Natural producer of K1 killer toxin – positive control for antifungal activity assays | National Collection of Yeast Cultures |
| *Metschnikowia pulcherrima* yAMV737 | Producer of a dark red iron-carrier – positive control for iron-siderophore production assays | This work – isolated from grapes |

**Supplementary Table 3.** Forward (fw) and reverse (rv) primers used in this study.

| **Primer** | **Sequence (5’🡪3’)** | **Use** | **Reference** |
| --- | --- | --- | --- |
| ITS1 (fw) | CTTGGTCATTTAGAGGAAGTAA | PacBio sequencing of ITS region for entire yeast collection | M. Gardes *et al.* (1993)^45^ |
| ITS4 (rv) | TCCTCCGCTTATTGATATGC |  | White *et al.* (1990)^46^ |
| P-ITS1 (fw) | TCCGTAGGTGAACCTGCGG | Sanger sequencing of ITS1 region for species identification of killer yeasts | J. Xie *et al.* (2008)^47^ |
| ITS2 (rv) | GCTGCGTTCTTCATCGATGC |  | M. Op de Beeck *et al.* (2014)^48^ |
| NL1 (fw) | GCATATCAATAAGCGGAGGAAAAG | Sanger sequencing of D1/D2 domain for species identification of killer yeasts | K. O’Donell *et al.* (1993)^49^ |
| NL4 (rv) | GGTCCGTGTTTCAAGACGG |  | K. O’Donell *et al.* (1993)^49^ |

**Supplementary Table 4.** Amplicon Sequence Variants obtained via PacBio sequencing the Internal Transcribed Spacer of all yeast isolates present in the collection pooled. Column B indicates the first hit after blasting sequences in column D. Column C indicates the amplicon length in base pairs. Species in red were discarded because there were no significant hits when blasting or they were regarded as contamination for being filamentous. *n/a:* not applicable.

**Supplementary Table 5.** Operational Taxonomic Units (OTUs) obtained after grouping the ASVs linked to same species. Columns B-G indicate the taxons phylum, order, class, family, genus and species of each OTU, respectively. Columnn H shows a representative sequence for each OTU chosen haphazardly.

**Supplementary Table 6.** Antagonistic yeast isolates classified by class of molecule likely produced responsible for the antifungal activity. In red those that lost the inhibitory phenotype whose class of compound produced could not be elucidated. *nd:* not determined*.*

**Supplementary Table 7.** Yeast isolates producing proteins with antifungal activity. Columns B-I show the mean and standard deviation of halo area measurements (in arbitrary units) against all eight target strains in the panel. Column J shows the phenotype cells present against the plant pathogen *B. cinerea*. Column K shows the primer used to send that yeast for species identification via Sanger sequencing. Columns L and M show the sequence identity (%) and query coverage (5), respectively. Column N shows the first hit when blasting and column O shows the sequence obtained after sequencing. *n/a*: not applicable (see **Supplementary Table 9**); *nd*: not determined.

**Supplementary Table 8.** Activity retained (%) of 100 – fold spent media of ten isolates against *C. tropicalis* under different pH and temperature conditions. The percentage values are normalized to 25 °C pH 4.

**Supplementary Table 9.** Values after halo measurment of screening of both cells and 200 - fold spent culture media of *A. pullulans* isolates against all eight target strains from the panel. Rows 2-8 show the mean of triplicates in arbitrary units and rows 10-16 contain those values calibrated into micafungin equivalents (µg/mL).

**Supplementary Table 10.** Set of 113 yeasts isolated from daily-consumed products. Column B shows the source type. Columns C-F indicate with 'yes' the antifungal activity against the corresponding target strains.

**Supplementary Table 11.** Halo area measurements of killer yeast candidates (**Supplementary Table 7)** calibrated into micafungin equivalents (µg/mL) against each target strains for every yeast isolate producing a high-molecular compound. *nd:* not determined; *n/a:* not applicable.

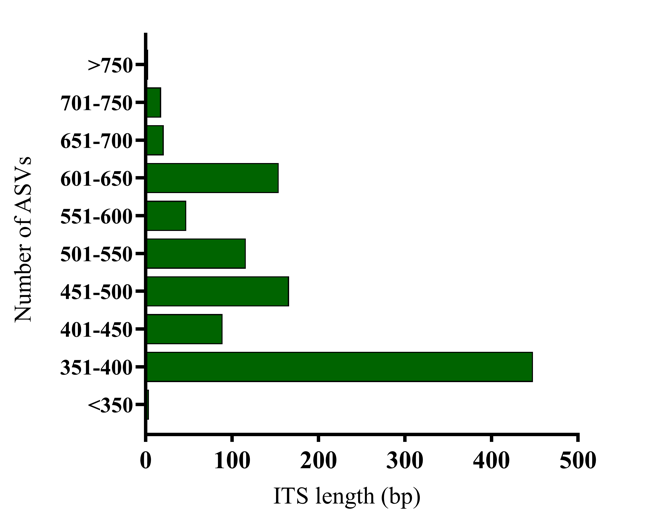

**Supplementary Figure 1.** Length distribution in base pairs of the Amplicon Sequence Variants (ASVs) obtained after sequencing the ITS region of the pooled yeast collection.

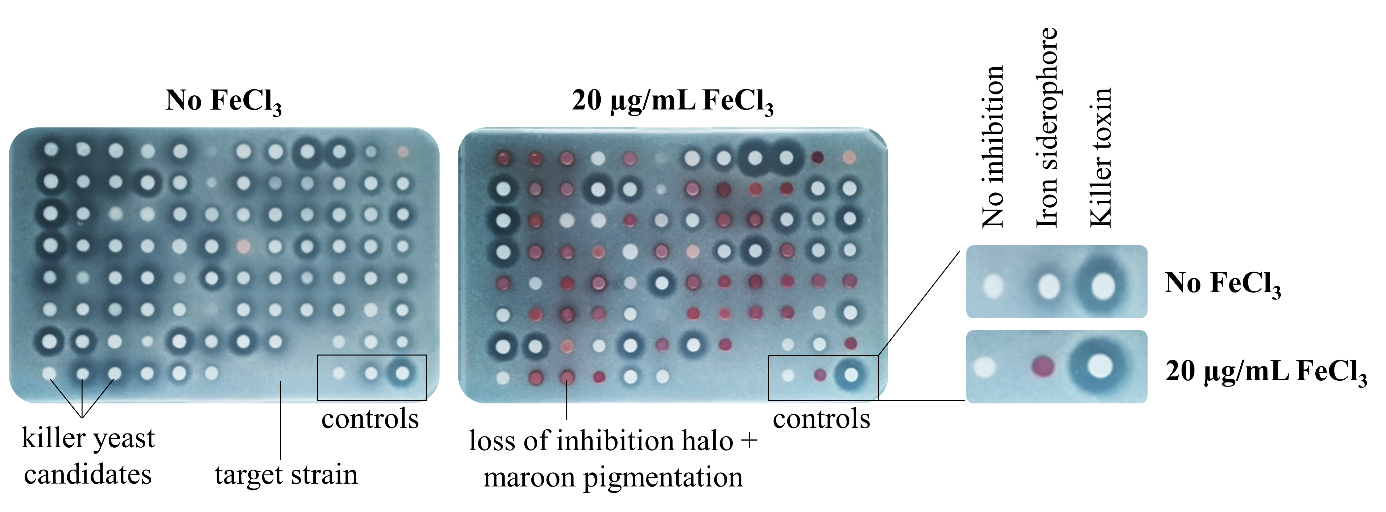

**Supplementary Figure 2.** Iron depletion screening assay**.** All yeasts that had shown an inhibitory halo in the primary screen were consolidated in three source plates. Isolates were spotted on top of an agar plate with trace amounts of iron (left) and with iron supplemented (right) using a manual pinner. The assay was performed in triplicate. The controls were from left to right, S. cerevisiae BY4741 (negative control – no inhibition), yAMV737 M. pulcherrima (pulcherriminic acid producer) and W. mrakii NCYC2251 (HM-1 toxin producer).

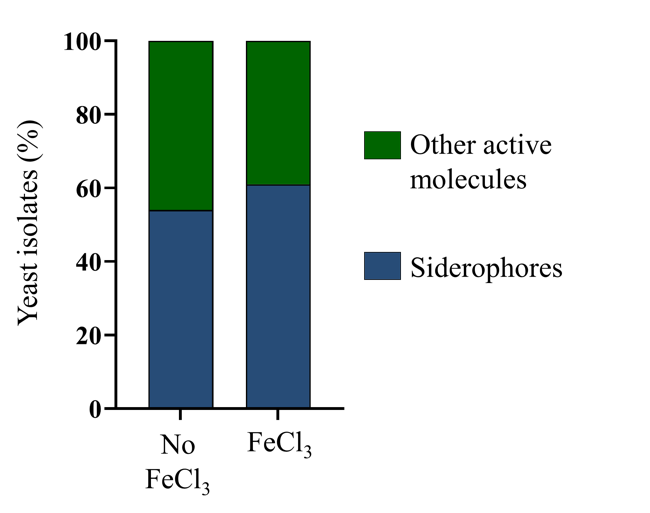

**Supplementary Figure 3.** Percentage of yeast isolates producing siderophores or other active molecules obtained when supplementing the enrichment media with additional iron.

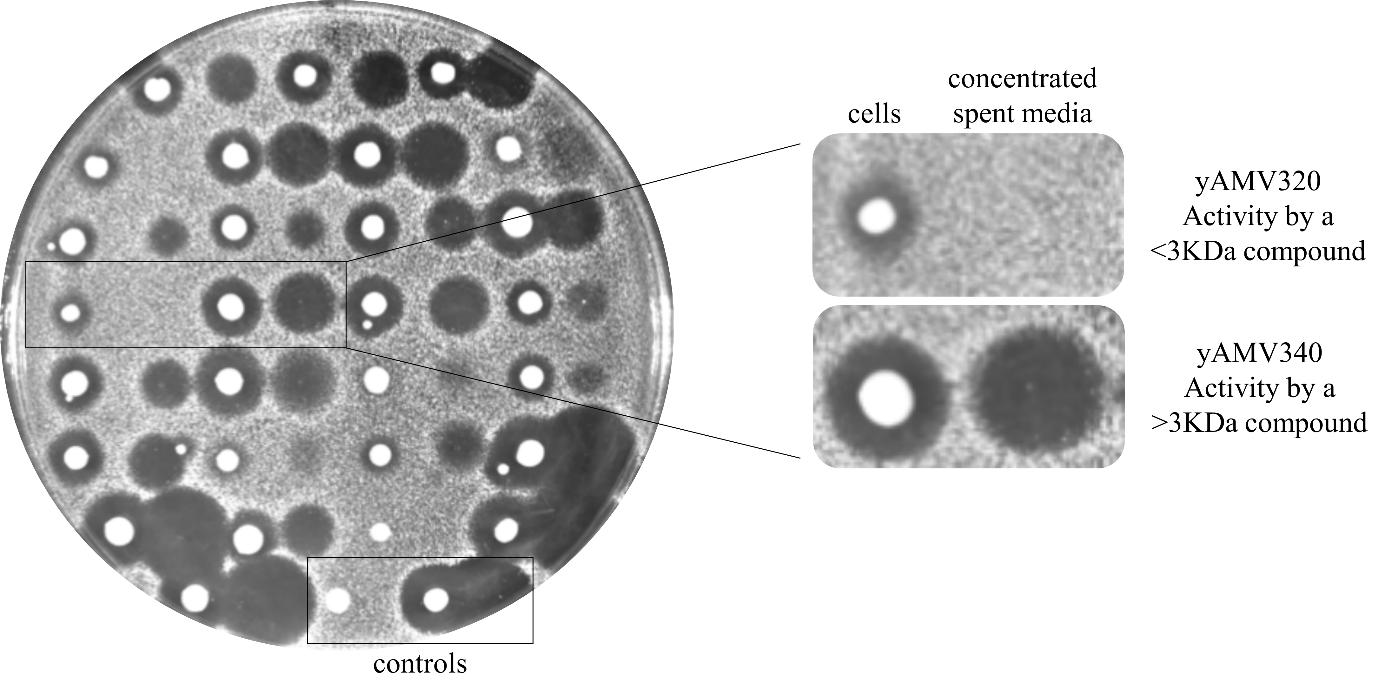

**Supplementary Figure 4.** Assay for protein content in the spent culture media**.** Cells and their concentrated spent media were tested against C. tropicalis. The spent media was concentrated using a 3KDa MWCO to discern yeast isolates producing proteins or other small compounds. The controls were S. cerevisiae BY4741 (negative control – no inhibition) on the left and W. mrakii NCYC2251 (HM-1 toxin producer) on the right.

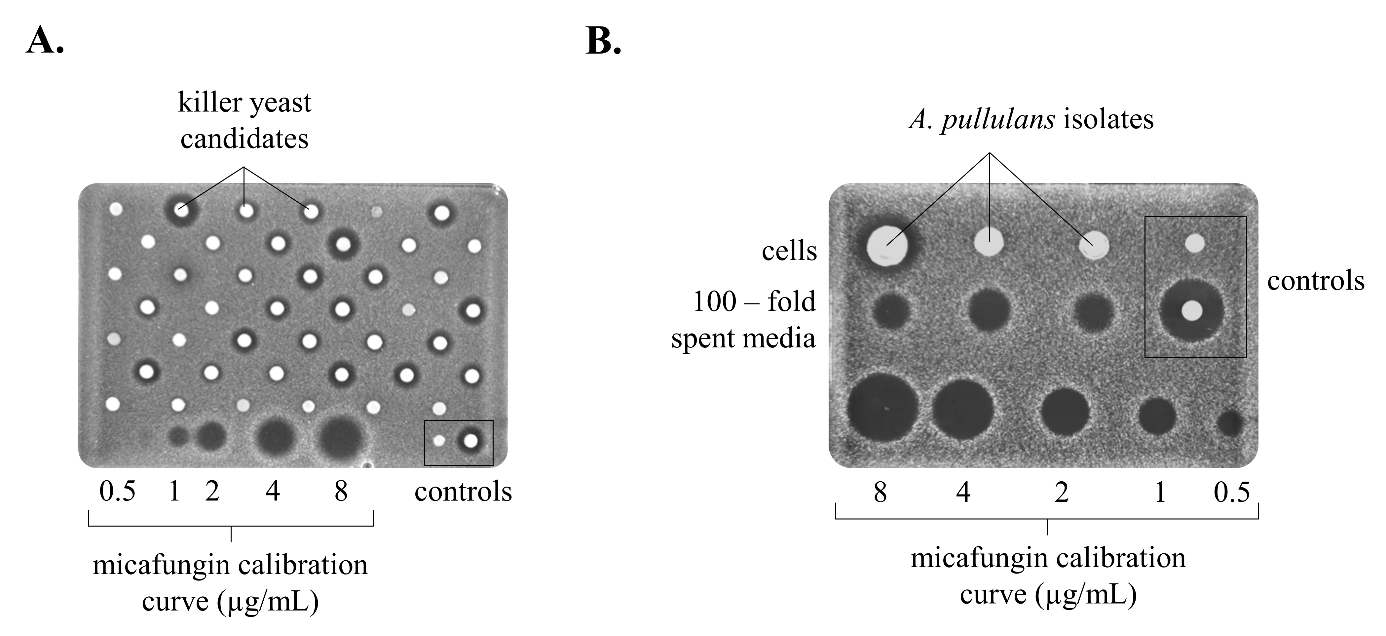

**Supplementary Figure 5.** Fungal inhibition assays for halo measurement and calibration with micafungin. **A.** Assays for killer yeast candidates using the manual pinner. The assay was performed in triplicate. **B.** Assay for A. pullulans isolates (from left to right, yAMV116, yAMV368 and yAMV598) both cells and 200 – fold spent media. The controls were S. cerevisiae BY4741 (negative control) and W. mrakii NCYC2251 (positive control). The assay was performed in triplicate.

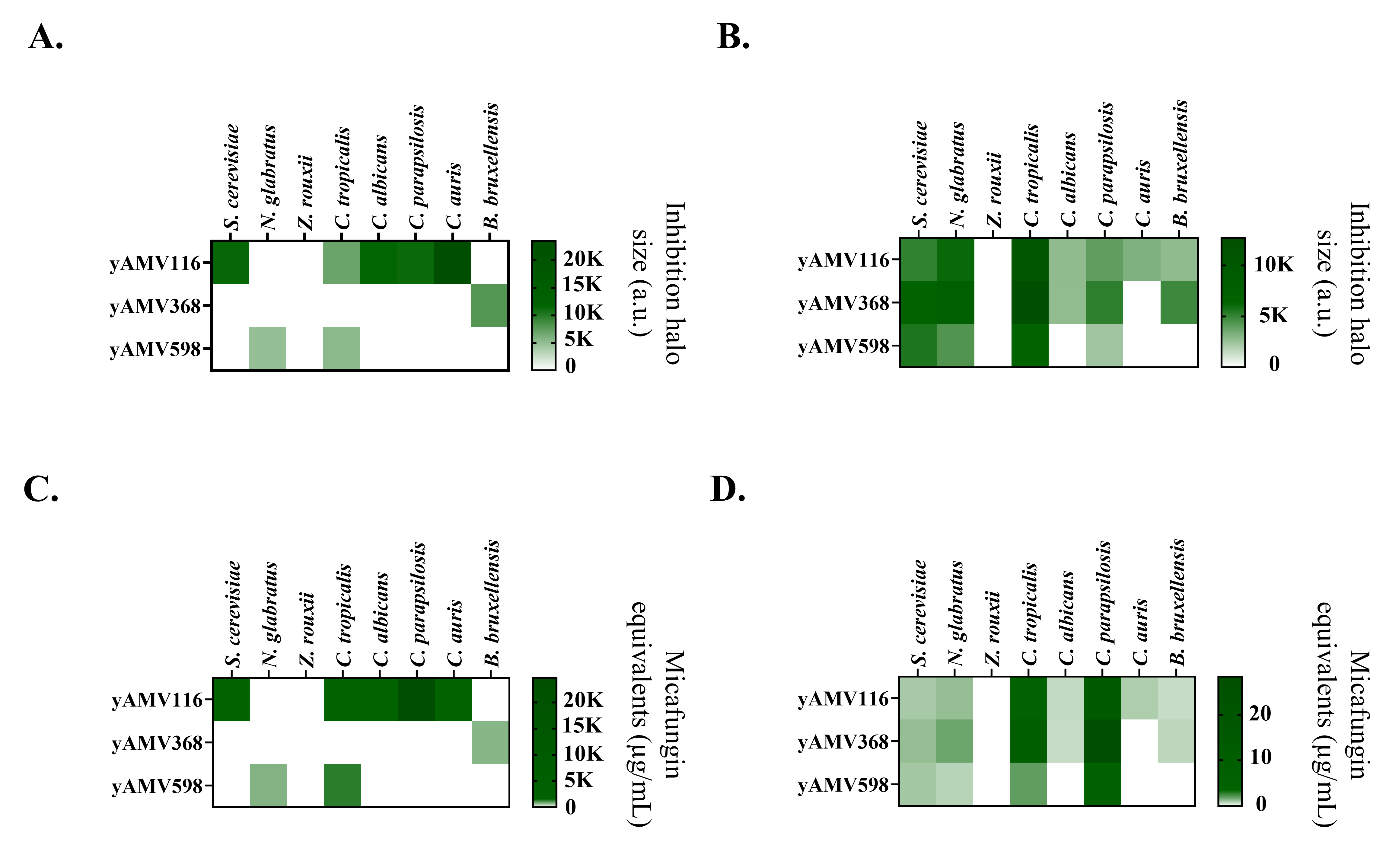

**Supplementary Figure 6.** Heatmaps depicting the halo size of A. pullulans yAMV116, yAMV368 and yAMV598 isolates against panel of eight target strains. **A.** Halo size in arbitrary units of cells, with their OD_600_ normalized to 50 units. **B.** Halo size of 200 – fold spent media. Using spent media refined the target range in comparison to using cells for the assay. **C.** Comparison of the halo sizes that were obtained when screening with cells to a calibration curve of the antifungal micafungin, units are given in “micafungin equivalent units”. **D.** Micafungin equivalents of 200 – fold spent media. The heatmaps depict the mean of triplicate experiments and the standard deviation is given in **Supplementary table 9.**

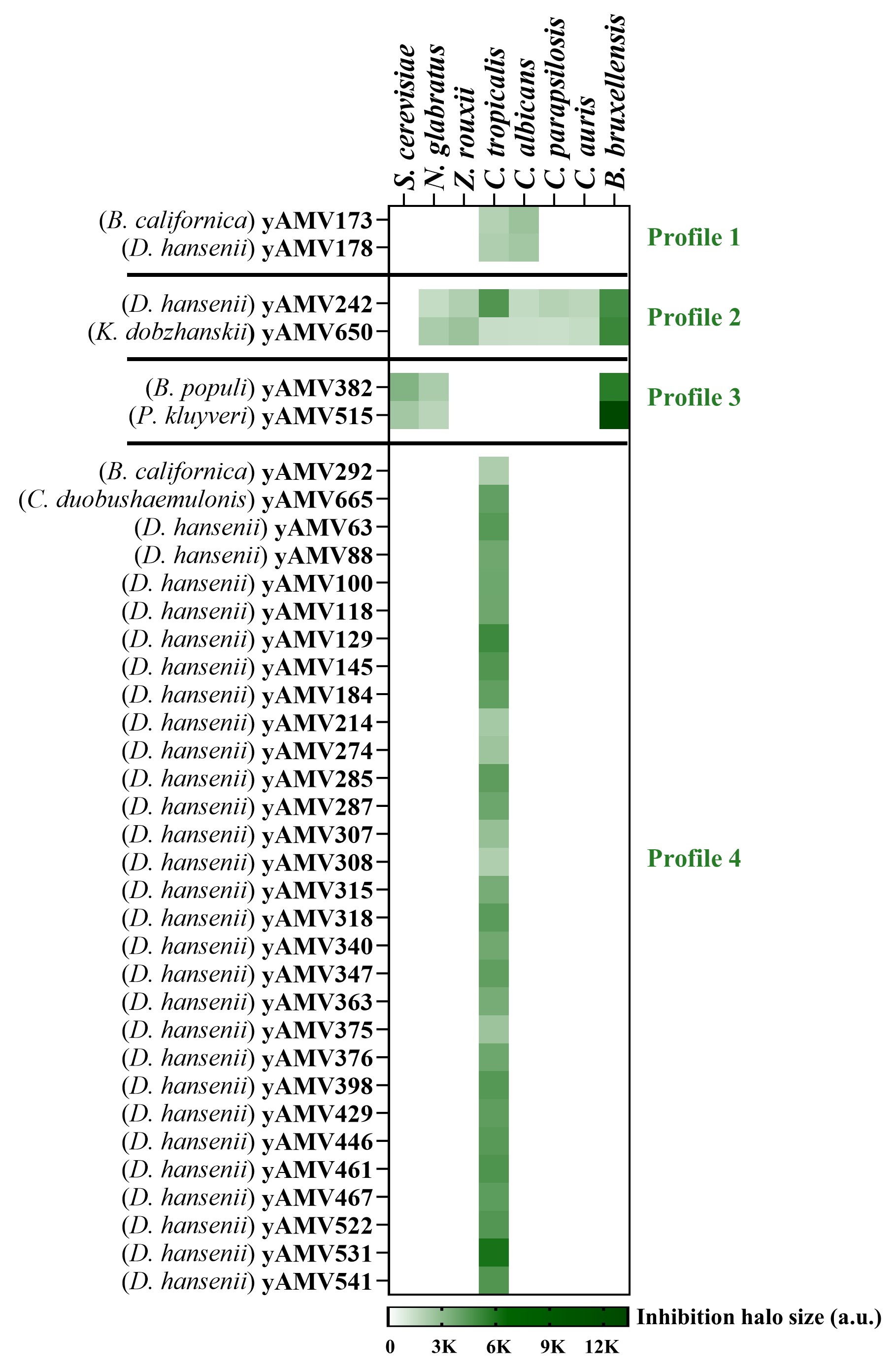

**Supplementary Figure 7.** Species-independent activity clusters of yeast isolates hinting possible shared binding receptors, targets or overall mechanisms of action.

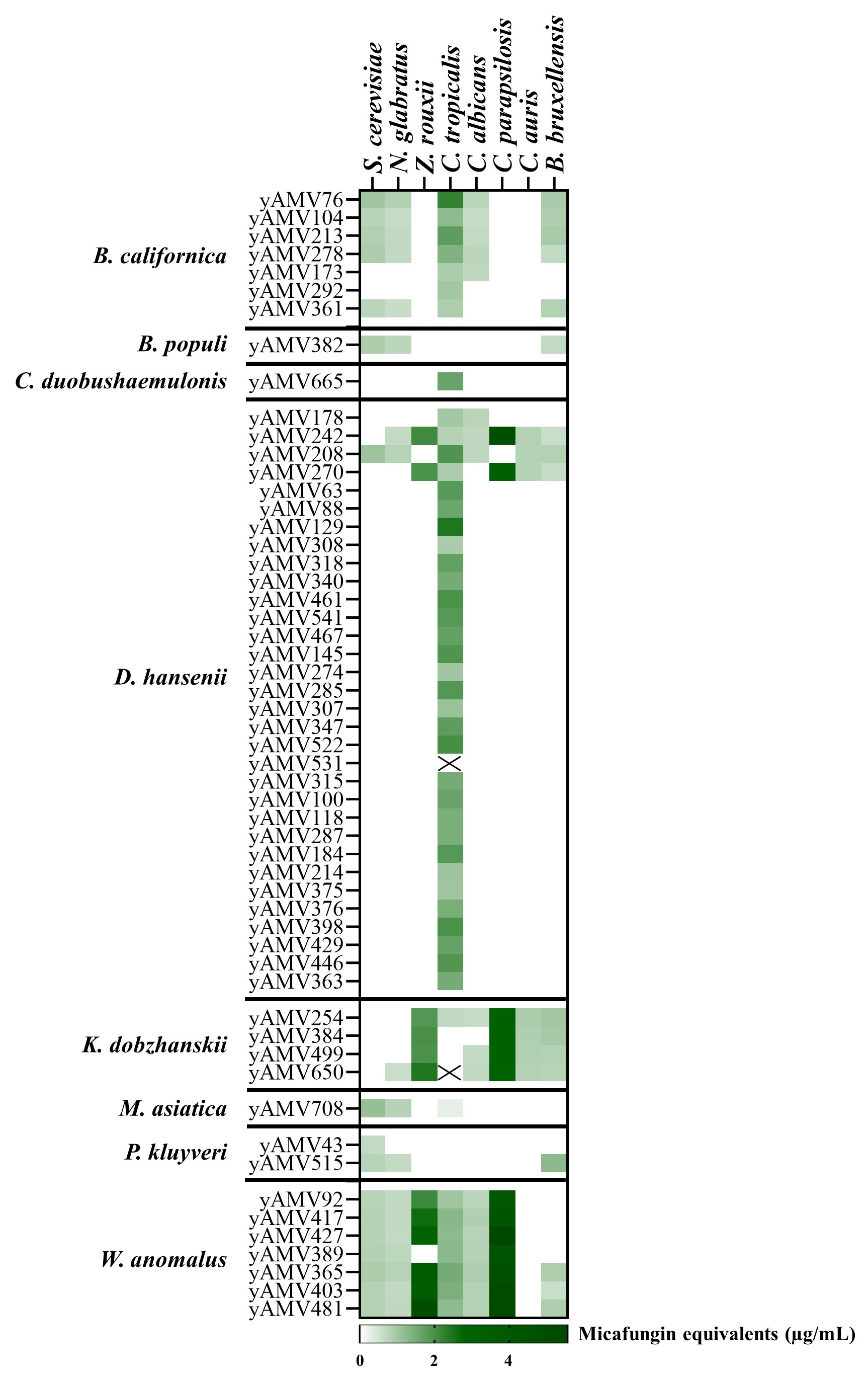

**Supplementary Figure 8.** Heatmap depicting inhibition halo size of yeast isolates calibrated to micafungin equivalents. Yeasts are divided by species. The cross (X) stands for data that were not determined. The heatmap depicts the mean of triplicate experiments and the standard deviation is given in **Supplementary Table 11.**
